# Massive Multiplexing of Spatially Resolved Single Neuron Projections with Axonal BARseq

**DOI:** 10.1101/2023.02.18.528865

**Authors:** Li Yuan, Xiaoyin Chen, Huiqing Zhan, Henry L. Gilbert, Anthony M. Zador

## Abstract

Neurons in the cortex are heterogeneous, sending diverse axonal projections to multiple brain regions. Unraveling the logic of these projections requires single-neuron resolution. Although a growing number of techniques have enabled high-throughput reconstruction, these techniques are typically limited to dozens or at most hundreds of neurons per brain, requiring that statistical analyses combine data from different specimens. Here we present axonal BARseq, a high-throughput approach based on reading out nucleic acid barcodes using *in situ* RNA sequencing, which enables analysis of even densely labeled neurons. As a proof of principle, we have mapped the long-range projections of >8,000 mouse primary auditory cortex neurons from a single brain. We identified major cell types based on projection targets and axonal trajectory. The large sample size enabled us to systematically quantify the projections of intratelencephalic (IT) neurons, and revealed that individual IT neurons project to different layers in an area-dependent fashion. Axonal BARseq is a powerful technique for studying the heterogeneity of single neuronal projections at high throughput within individual brains.

## Introduction

The mouse brain contains over 70 million neurons (Herculano-Houzel et al., 2006), and the combined length of their axonal trees stretches thousands of kilometers; in the human brain, there are orders of magnitude more. These axons form the scaffolding for neural circuits and hence for computation. Tracing these projections represents a formidable challenge. Traditionally, there are two main approaches. At one extreme, the projections of single neurons can be reconstructed at high resolution by labeling neurons one at a time, using e.g. the Golgi method or more modern sparse labeling based on viral delivery of fluorophores such as green fluorescent protein (GFP). Such single-neuron methods have undergone impressive advances in recent years, but even today allow the multiplexing of at most dozens of neurons from a single brain region (Winnubst et al., 2019; Peng et al., 2021; Gao et al., 2022; Qiu et al., 2024). Alternatively, the projections of major projection pathways can be assessed using bulk tracing methods. For example, a bolus of virus expressing a fluorophore can be injected into one brain area, enabling the major projections of neurons within that area to be visualized by microscopy. These techniques have been used to systematically map the mesoscopic projections of the mouse brain (Oh et al., 2014; Harris et al., 2019; Muñoz-Castañeda et al., 2021). Bulk methods reveal the projections of large populations of neurons, but at the cost of single-cell resolution. Thus, there is a tradeoff between throughput and single-cell resolution in traditional methods.

We have recently developed a novel suite of nucleic acid barcode-based tracing techniques, which provide a third alternative. The first-generation method for exploiting barcodes in the context of circuit mapping was Multiplexed Analysis of Projections by Sequencing (MAPseq, Kebschull et al., 2016a). MAPseq can reliably and simultaneously map the projections of hundreds of thousands of individual neurons in a single experiment. MAPseq uniquely labels individual neurons by introducing random RNA sequences (“barcodes”) via infection with a barcoded viral library. These random barcodes fill the cells and are co-expressed with a protein that has been engineered to bind to the barcode and drag it to distant axonal terminals. The pool of unique barcode identifiers is effectively infinite; even a 30 nucleotide (nt)-sequence has a potential diversity of 4^30^≈10^18^ barcodes, far surpassing the ∼10^8^ neurons in the mouse brain. This high diversity implies that most neurons are uniquely labeled. The barcode RNA can then be extracted from the axons in an area of interest to determine which neurons project there; the number of molecules with a specific barcode sequence collected from a region is used as a proxy for the strength of the projection (i.e., axonal volume) of that particular barcoded neuron, in much the same way that GFP intensity is used as a proxy for projection strength in conventional bulk injections (Kebschull et al., 2016a). Because high-throughput sequencing can quickly and inexpensively distinguish these barcodes, MAPseq can uncover the projections of hundreds of thousands of individual neurons in parallel within a single brain (Huang et al., 2020; Chen et al., 2022). The throughput of MAPseq for assessing single neuron projection patterns in a single brain is currently unmatched by any other approach.

MAPseq was the first approach to exploit barcoding for neuronal mapping. However, because it relies on bulk sequencing of homogenized tissue, its spatial resolution is determined by the precision of dissection. To achieve higher resolution, we developed BARseq (Barcoded Anatomy Resolved by Sequencing), the next generation of sequencing-based tracing (Chen et al., 2019; Sun et al., 2021). BARseq relies on *in situ* sequencing. Unlike conventional *in situ* hybridization, which uses a complementary probe to detect a specific RNA molecule in the cell, *in situ* sequencing obtains the exact sequence of each RNA target. This is a key difference, as the RNA barcode in any given cell is unique, unknown and highly diverse, making it very challenging to design probes in sufficient numbers for the desired targets. In contrast, *in situ* sequencing makes it straightforward to discriminate an almost infinite number of sequences. Combining BARseq-based sequencing of somatic barcodes and endogenous gene expression with MAPseq-based dissection and sequencing of barcodes in the axons allows us to associate the projection patterns of individual neurons with soma locations in a highly multiplexed manner (Chen et al., 2019; Sun et al., 2021). However, because spatial resolution in MAPseq is limited by the dissection of brain areas prior to bulk sequencing, axonal projection patterns in this MAPseq/BARseq combined approach can only be crudely resolved. MAPseq and BARseq have been repeatedly validated using multiple methods in a wide range of brain areas (Kebschull et al., 2016a; Han et al., 2018; Chen et al., 2019, 2022; Gergues et al., 2020; Huang et al., 2020; Klingler et al., 2021; Mathis et al., 2021; Muñoz-Castañeda et al., 2021; Sun et al., 2021; Hausmann et al., 2022; Tsoi et al., 2022; Webb et al., 2022; Zeisler et al., 2023). In particular, we demonstrated that barcode transport is uniform over long distances (>10 mm; Kebschull et al., 2016a; Han et al., 2018; Chen et al., 2019, 2022; Gergues et al., 2020; Huang et al., 2020; Klingler et al., 2021; Mathis et al., 2021; Muñoz-Castañeda et al., 2021; Sun et al., 2021; Hausmann et al., 2022; Tsoi et al., 2022; Webb et al., 2022; Zeisler et al., 2023).

We therefore set out to increase the spatial resolution with which highly multiplexed axonal trajectories can be resolved using *in situ* sequencing. To achieve this, we developed a method, “axonal BARseq,” for sequencing individual axonal “rolling circle colonies”, or “rolonies”, *in situ*. Axonal BARseq allows much finer resolution of the spatial organization of axonal projections than can be achieved with MAPseq. Using this approach, we identify the projections of thousands of individual axons projecting from a single localized injection in a single mouse, increasing throughput beyond current methods and eliminating the need to register injections across samples. Axonal BARseq has the potential to scale up to multiple injection sites and reveal projections from multiple sites, raising the possibility of sampling brain-wide projections from multiple neuronal populations at single-cell resolution.

## Results

Here we describe axonal BARseq, a highly multiplexed method for reconstructing axonal trajectories. We first describe the optimizations necessary to achieve single molecule sequencing of barcodes in axons. Next, we demonstrate its application to determine axonal projections from auditory cortex. We confirm that the single-neuron projection patterns obtained by this method are consistent with previous single-neuron approaches. We then show that the high resolution and multiplexing of axonal BARseq reveals the statistical structure of single neuron projections to different laminae in different areas.

### Optimizing BARseq to achieve axonal resolution

We have previously demonstrated *in situ* read-out of barcodes expressed in somata (Chen et al., 2019; Sun et al., 2021). This is a much easier problem than the present challenge of reading out single axonal barcodes, because there are several orders of magnitude more barcodes in somata (10^3^-10^4^; Kebschull et al., 2016a). We therefore sought to maximize the sensitivity of *in situ* read-out of barcodes to achieve high-efficiency single-barcode readout of barcodes transported millimeters or centimeters from their soma of origin.

To increase the sensitivity of *in situ* sequencing of axonal barcodes, we modified the sequencing protocols originally developed for barcodes in somata (Chen et al., 2018, 2019). The basic *in situ* sequencing protocol consists of (1) injection with a Sindbis virus engineered to express a diverse barcode library; (2) tissue preparation 24-48 hours after infection; (3) preparation of rolonies (nanoballs of DNA generated by reverse transcription of the RNA barcode, followed by gap-filling padlock-extension, ligation, and rolling circle amplification) in thin brain slices; (4) *in situ* sequencing by synthesis using standard Illumina reagents: sequential four-color imaging of each base in the barcode of each rolony (see *Methods*; Fig. 1A, SupFig. 1A). We optimized the reverse transcriptase used and the gap-filling procedure (SupFig. 1B-D), which increased the sensitivity of barcode detection to an efficiency of 20.2% compared to RNA *in situ* hybridization (SupFig. 1E-F). In addition, we engineered a Sindbis virus using a second-generation carrier protein (VAMP2nλ), which carried barcodes more efficiently than our previously described carrier protein (see *Methods*).

**Fig. 1.**
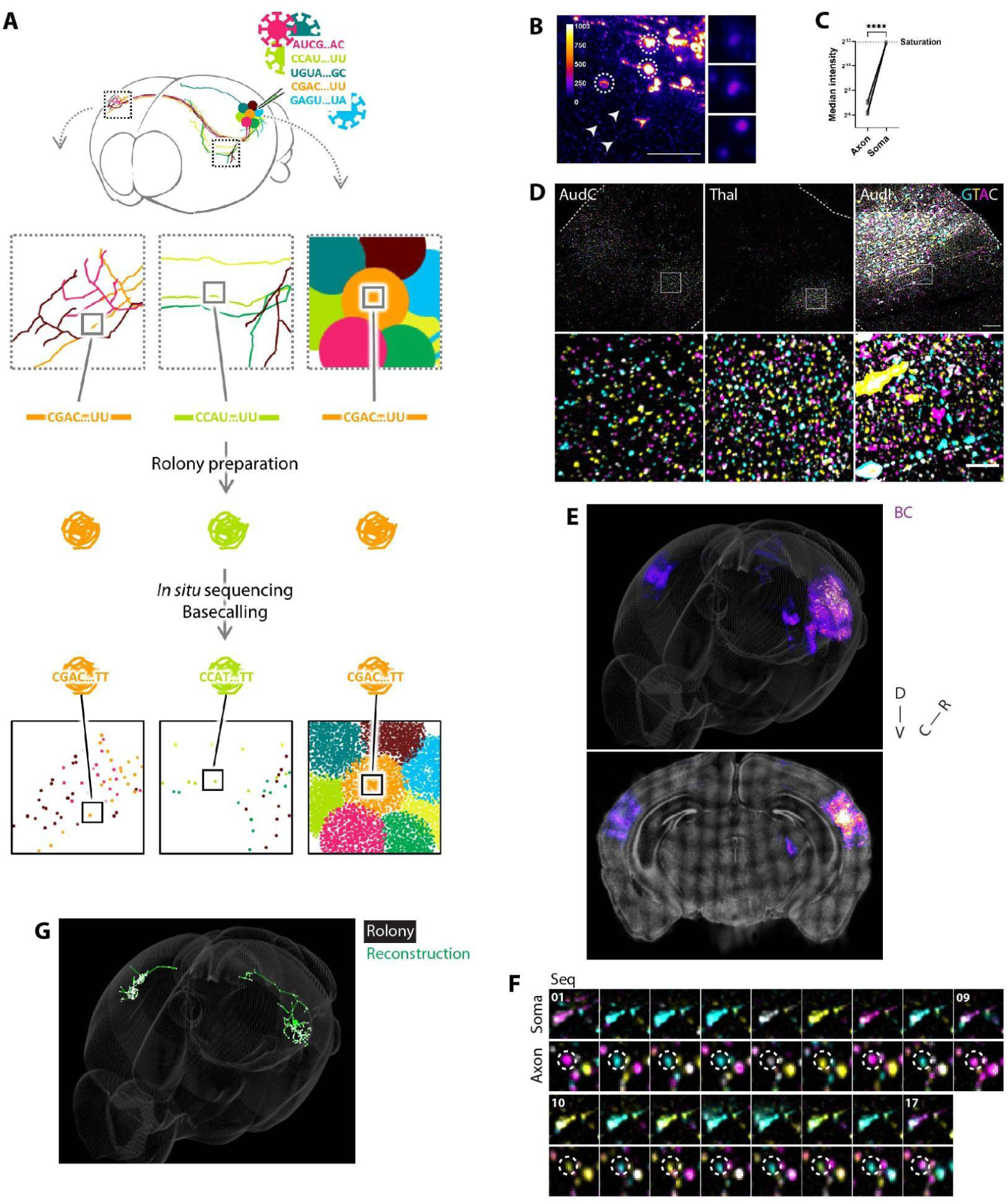
Overview of axonal BARseq. (**A**) Workflow. The brain is injected with barcoded viral library. After 24-48 hrs of expression, during which RNA barcodes are transported to axon terminals, where they are amplified into rolonies and sequenced. (**B-C**) Single rolonies in axons have significantly weaker signals compared to somatic rolonies. (**B**) Representative image of somatic and axonal rolonies; dotted circle: somatic rolonies; arrow: axonal rolonies, with zoom-in views shown on the right; rolony intensity is color coded. Scale bar: 100 µm. (**C**) Quantification of intensity between axonal and somatic rolonies. Due to the large intensity difference between somatic and axonal rolonies, proper exposure for axonal rolonies often results in saturation of somatic rolonies. Paired *t*-test. (**D**) Representative images of axonal and somatic rolonies in AudC/I and ipsilateral thalamus. Images are from the first cycle of *in situ* sequencing. Dotted line, anatomical boundaries. Scale bar: top, 100 µm; bottom, 25 µm. **(E**) Registered barcode signals in CCFv3. Top, data in 3D model. Gray, brain outline. Bottom, coronal view of 25 µm of the sample. Gray, DAPI. (**F**) Representative images of *in situ* sequencing soma and a single axonal rolony with the same barcode. Soma ROI, 30.25 µm × 30.25 µm from injection site; axonal rolony ROI, 14.85 µm × 14.85 µm from ipsilateral thalamus. In total, 17 sequencing cycles are shown. (**G**) An example of tracing tracks for a single barcoded neuron reconstructed by connecting rolonies. Rolony location is indicated in white; soma location is indicated as a large green dot in ipsilateral cortex. AudC/I, contra/ipsilateral auditory cortex; Thal, thalamus; BC, barcode; D, dorsal; V, ventral; C, caudal; R, rostral; Seq, sequencing cycle.

A further challenge of single-barcode axonal sequencing is to achieve the requisite sensitivity and accuracy during successive rounds of *in situ* sequencing. The signal from a single axonal rolony is not as bright as that from larger somata because somata contain many copies of the same barcode (Fig. 1B-C). In addition, alignment of single rolonies across successive rounds of imaging poses additional challenges compared with alignment of somata. Overcoming these challenges required considerable modifications and optimization (see *Methods*, SupFig. 2-4).

**Fig. 2.**
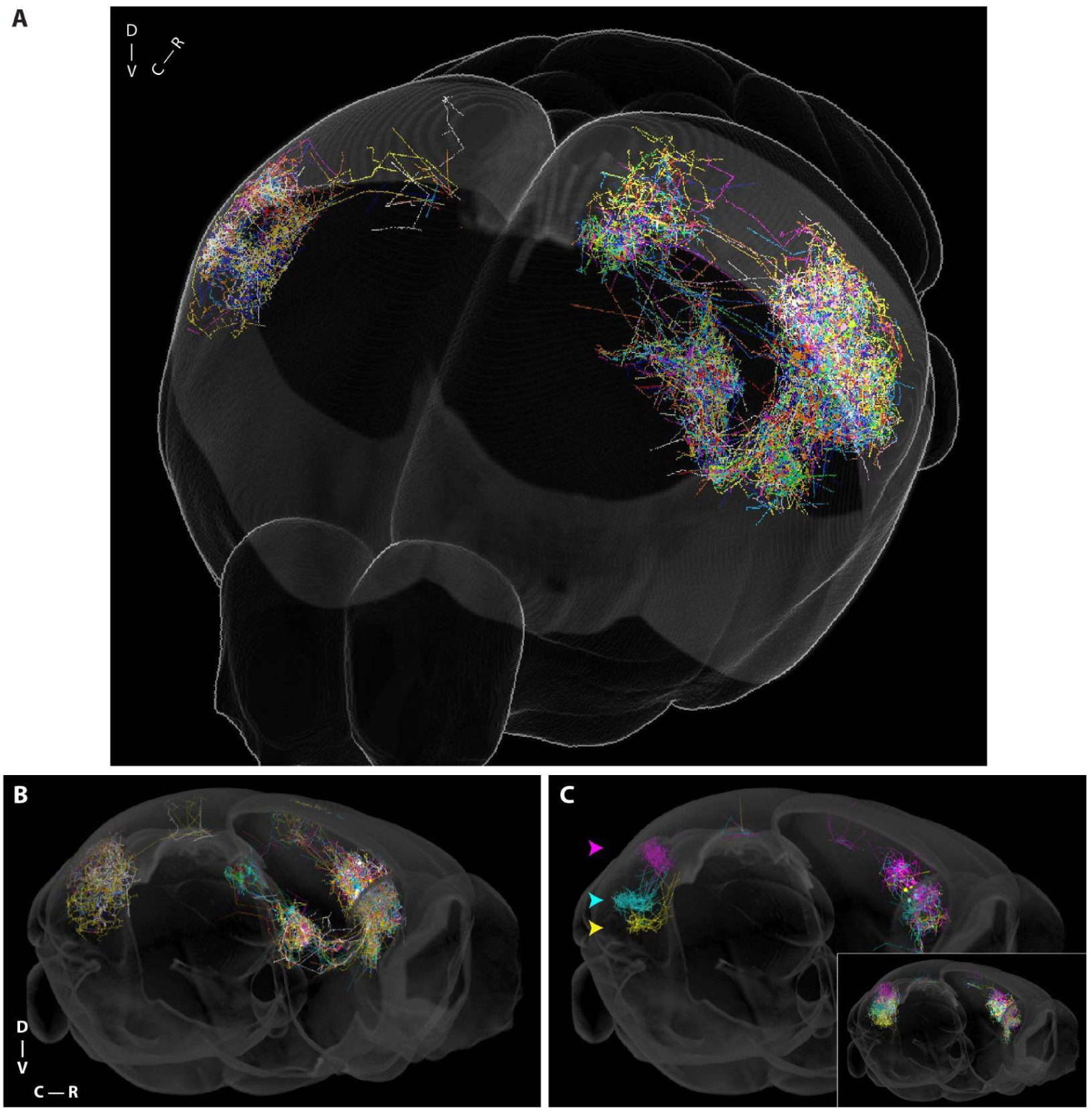
Axonal barcodes can be used to reconstruct axonal projection in anterograde and simulated retrograde tracing. (**A-B**) Single-cell reconstruction from 100 barcoded neurons, 25 from each major cell type (ET, CT, ITc, ITi) as described below. Somas (colors randomly assigned) are indicated by large dots in the left hemisphere. Not all tracts were imaged. Note that these reconstructions are approximate because many fine processes are not detected, and that only the 2 mm *shaded region* was reconstructed (the entire brain is included for reference). The reconstructions are used for visualization purposes only; all quantifications rely directly on the rolonies themselves rather than the reconstructions. (**C**) Identifying cells with focal projections to the same locations by simulated retrograde tracing. Cells from three simulated retrograde injections are indicated by arrows in the right hemisphere. (*inset)* For each simulated injection, 25 neurons were randomly selected and plotted, regardless of whether their projections were focal or broad.

**Fig. 3.**
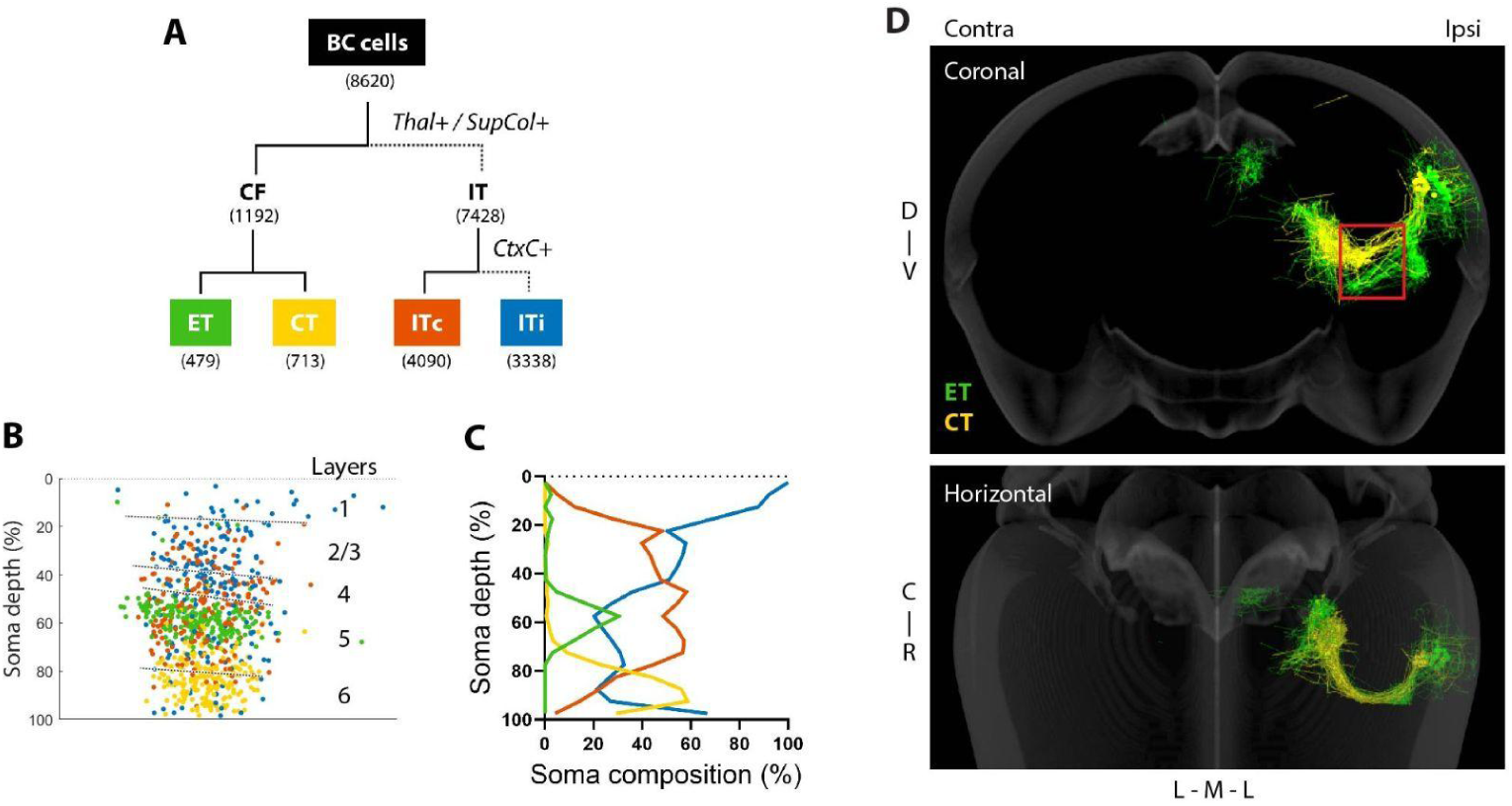
Cells can be divided into major cell types using projection targets and axonal trajectory. (**A**) Barcoded cells were divided into four cell types: ET, CT, ITc and ITi. Cells were divided into CF/IT using Thal/SupCol projection; IT cells were divided into ITi/ITc using contralateral cortical projection. SupCol was identified using the midbrain area in CCFv3. Solid line, with the specific projection; dotted line, without the projection. Numbers next to the group names indicate cell counts of the groups. (**B-C**) Soma depth of four cell types in the injection site. (**B**) Scatter plot of 200 randomly selected cells per group. Layer annotations are from CCFv3. (**C**) The distribution of the four cell types along depth. Cell types were color coded as in Fig. 3A. (**D**) Projection tracts of CT and ET neurons consistent with spatial features of both cell types, 60 randomly selected neurons per type. Red box, region of interest for CT/ET grouping. Coronal view, top; horizontal view, bottom. CT/ET rolonies with region boundaries were shown in SupFig. 7F-G. SupCol, superior colliculus; CtxC/I, contra/ipsilateral cortex; CF, corticofugal; IT, intratelencephalic; ET, extratelencephalic; CT, corticothalamic; L, lateral; M, medial.

**Fig. 4.**
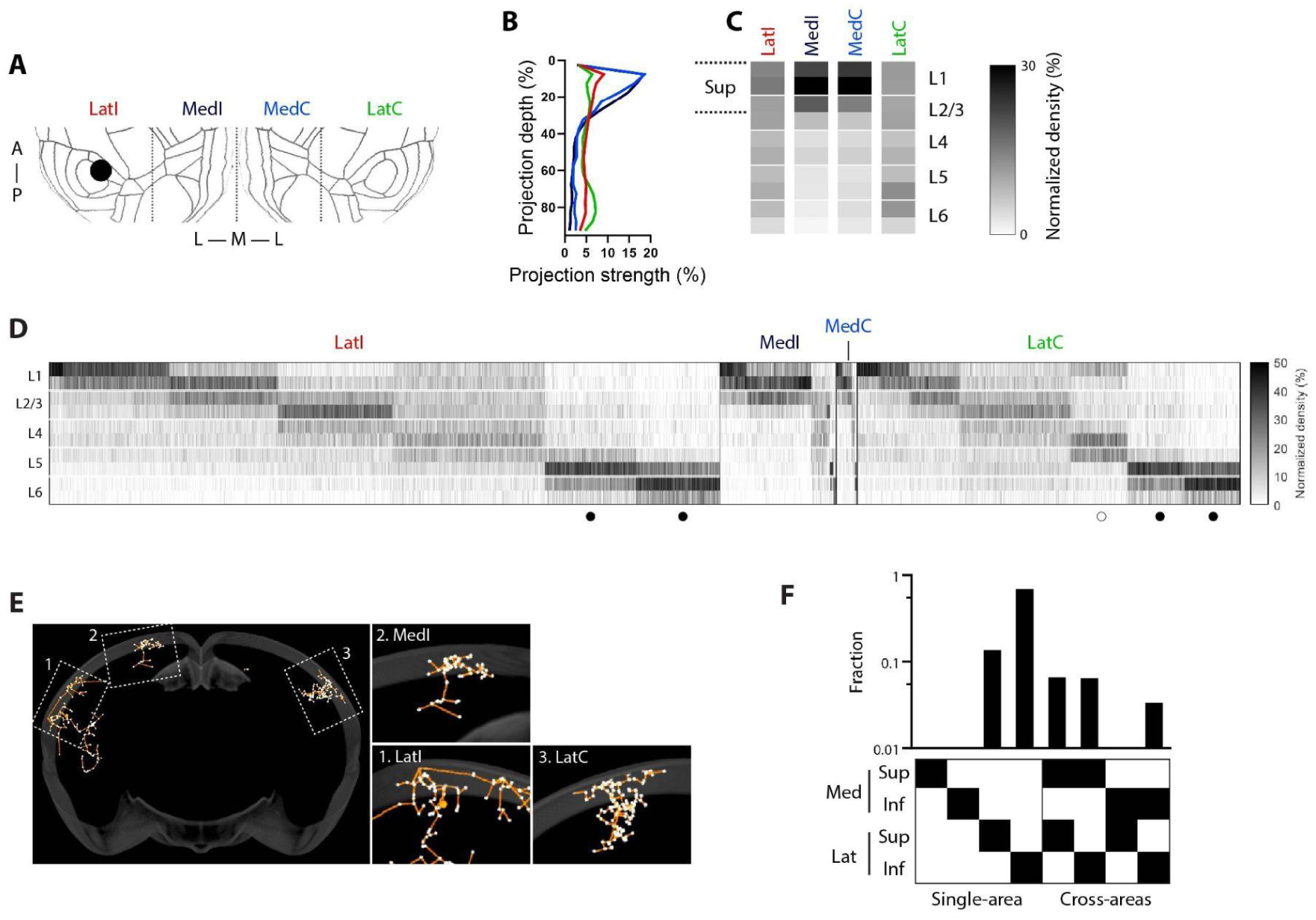
IT projections demonstrate diversity across cortical areas, within individual areas, and across different areas within single cells. (**A**) Division of cortex into medial and lateral targets on flatmap: LatI, MedI, MedC, and LatC. Black disk, injection site. (**B-C**) Different laminar patterns of bulk projections in medial and lateral cortical targets. (B) Frequency distribution of projection strength along projection depth. Bin size, 5% depth. (C) Heatmap of the normalized projection density in each layer. Each layer is divided into upper and lower halves. Superficial layer (Sup) refers to projections from L1 to upper L2/3. To calculate the normalized density, rolony counts were normalized to the layer thickness and the total rolony counts per column. Cell counts: LatI, 6,878; MedI, 1,196; MedC, 212; LatC, 3,931. (**D**) Distinct laminar patterns of single-cell projections in medial and lateral cortical targets. The heatmap illustrates the normalized projection density in each layer, with one cell per column. Laminar patterns were clustered using *k*-means. (**E**) Coronal view of representative single-cell reconstruction (orange) and axonal rolonies (white) of an IT neuron with multiple cortical targets. Soma, big orange dot in AudI. Zoom-in view: 1, LatI; 2, MedI; 3, LatC. (**F**) Sup/Inf projection probability depends on Lat/Med projection patterns. Within a given area, neurons are categorized based on their predominant projection type (x-axis), e,g, a neuron is classified as ‘Sup’ if > 50% of its projections are superficial; otherwise, it is classified as ‘Inf’. Neurons that project only to a single area project only to Lat, and predominantly inferiorly, whereas neurons that project to both Lat and Med project with nearly equal probability to Sup and Inf. LatI/C, ipsi/contralateral lateral; MedI/C, ipsi/contralateral medial; A, anterior; P, posterior.

### Axonal BARseq of projections from auditory cortex

To assess the utility of these optimizations, we used axonal BARseq to reconstruct projections from mouse primary auditory cortex. Two days following unilateral viral injection, we performed 17 cycles of sequencing of coronal sections centered +/- 1 mm around the injection site (108 serial 20 µm sections). These sections contained many of the main projection targets of the auditory cortex, including most of the contra- and ipsilateral auditory cortices (AudC, AudI), contra and ipsilateral visual cortex (VisC, VisI), ipsilateral thalamus (Thal), part of the ipsilateral striatum (Str) and part of the ipsilateral superior colliculus (SupCol) (Fig. 1D-E, SupFig. 5A, SupTable 1). A total of 8,620 unique axonal barcodes (obtained from 492,950 sequenced rolonies) were used for analysis, with a median of 40 rolonies per barcode. About half (3,698/8,620) of the reliably detected axonal barcodes could be associated with somata whose position could be confidently determined (Fig. 1F, SupFig. 5B); the remaining barcodes could not be precisely localized due to various experimental and analytical challenges (see *Methods*). However, for most of the subsequent analyses (with the exception of Fig. 5 and partial SupFig. 9), we used the entire set of axonal barcodes as the analyses do not require information about soma depth. Rolonies close to somata at the injection site were excluded (see *Methods*). Barcode statistics are summarized in SupFig. 5C-G; for details of manual validation of sensitivity and accuracy see *Methods*.

**Fig. 5.**
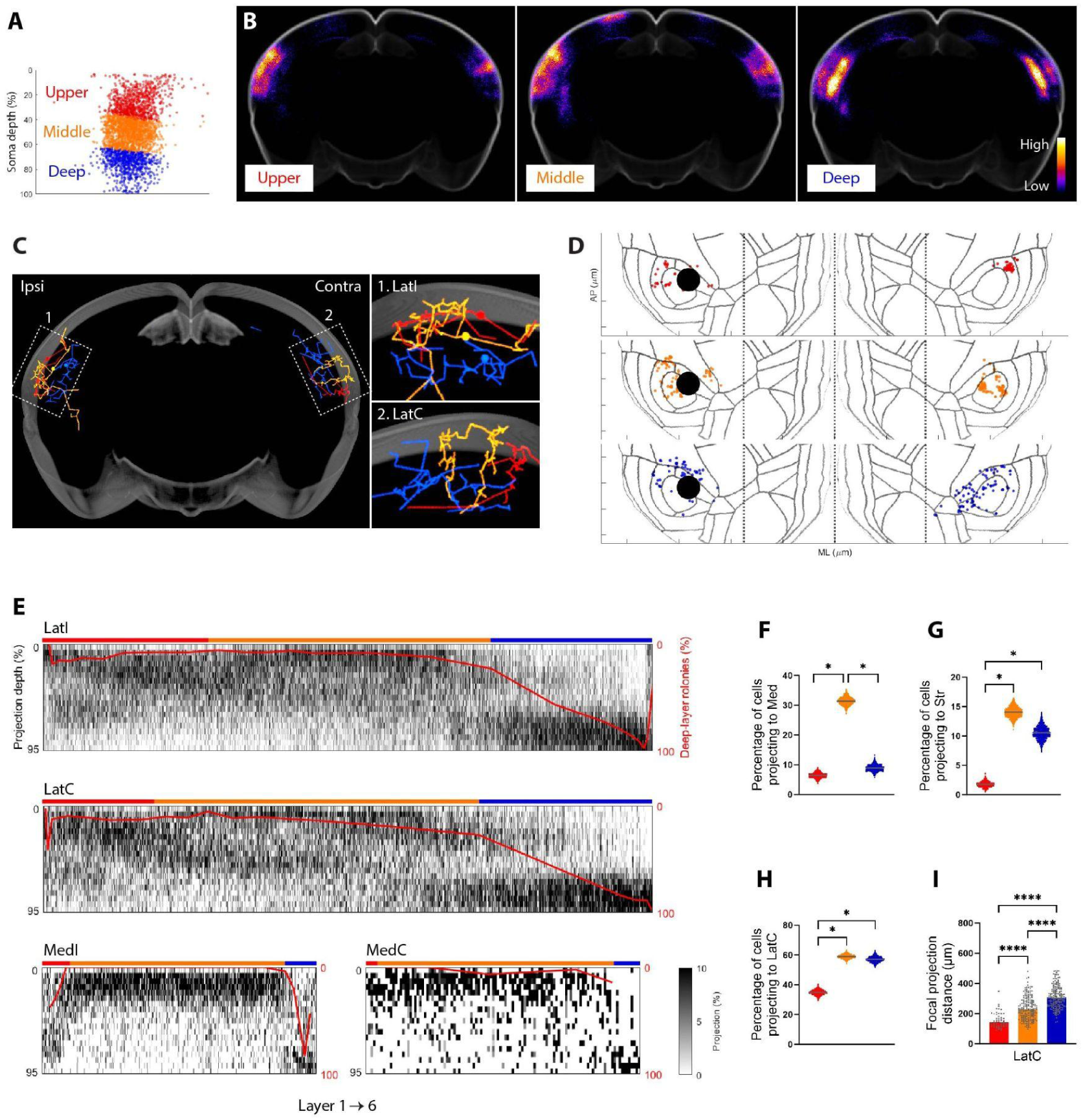
Diversity in laminar terminations, morphology, and targets across three groups of IT cells. (**A**) Lat-projecting IT cells were divided into three groups using soma layers: upper, layer 1 and 2/3; middle, layer 4 to upper layer 5; deep, lower layer 5 and layer 6. Cell counts: upper, 853; middle, 1,450; deep, 806. (**B**) The three groups have different rolony distribution patterns at bulk level. 806 barcodes per group were randomly selected for display purposes. Images shown are sum projections of coronal views. (**C**) Representative single-cell reconstructions of three biLat cells. One cell for each group. Zoom-in panel: 1, LatI; 2, LatC. Soma, big dots in AudI. (**D**) Rolony location of individual neurons in Fig. 5C on cortical flatmap. Dotted lines, Med-Lat boundaries. Ticks on x/y-axis, 2000/1000 µm; x/y-axis ratio, 1:1. (**E**) At the single-cell level, deep somata mainly project to deep layers, while upper and middle somata preferentially project to upper-middle layers. Heatmap, frequency distribution of rolony along depth; bin size, 5% depth; one cell per column. Cells were sorted by layer location of somata (x-axis). Red line: percentage of deep-projection (lower layer 5 and layer 6). Cells were binned into 5% bins using soma depth, and median percentage of deep-projection was calculated per bin. Cell counts: LatI, 3,012; MedI, 525; MedC, 113; LatC, 1,612. (**F-H**) Middle somata have a higher percentage of cells that project to the medial cortical areas (F), while upper somata have a lower percentage of cells projecting to the ipsilateral striatum and contralateral lateral areas (G-H). Gray line, median; *, significant difference with no overlap CI. (**I**) Upper/middle somata have more focal projection in LatC compared to deep somata. The focal projection distances were computed based on distances between rolonies (see *Methods*). Bar graph, median; dots, individual cells. Cell counts: upper 54; middle, 186; deep, 178. Kruskal-Wallis test, Dunn’s test for multiple comparisons. AP, anterior-posterior axis; ML, medial-lateral axis.

In conventional neuroanatomical single-neuron reconstructions, tracing requires that neuronal processes be filled with markers such as GFP or dyes, enabling visualization of axons as continuous structures. Disruption of this continuity due to errors in sample preparation or imaging can disrupt tracing and lead to catastrophic errors in reconstruction, potentially causing misattribution of an axon to the incorrect soma of origin. By contrast, because BARseq assigns axons to their soma of origin on the basis of their barcode sequences, assignments can be accurate even when barcodes are sparse. Errors in appropriate attribution of a barcode (e.g. due to sequencing errors) are rare and, importantly, are not catastrophic because they are independent, i.e. a given error affects only a single rolony.

We identified 8,620 axonal barcodes outside the injection site, including ipsi- and contralateral cortex (CtxI and CtxC), thalamus, caudal striatum, and superior colliculus (SupFig. 6A). For visualization purposes, it can be convenient to connect barcode rolonies to generate images that are similar to conventional neuroanatomical reconstructions. An example of such a connect-the-dots visualization, with straight lines linking nearby barcodes with the same sequence (see *Methods*), is shown in Fig. 1G. In cases where the inter-barcode distance is large, this reconstruction is only an approximation, since axons can sometimes take tortuous paths, which may not be fully captured by this approximation. However, these reconstructions are used for display purposes only; all quantifications rely directly on the rolonies themselves rather than the reconstructions. Fig. 2A-B shows the trajectories of 100 neurons (out of 8,620), color-coded for display purposes (see 3D rotation animation, SupVideo).

The large number of barcoded single neurons allowed us to identify subpopulations of neurons with distinct projection patterns. Fig. 2C (*inset*) shows a simulated contralateral retrograde injection of three colors. Among these neurons, subsets could be identified that projected very narrowly to specific patches (Fig. 2C). The identification of such subpopulations is facilitated by the high density of labeling obtained with axonal BARseq and would have been difficult to identify using conventional anterograde or retrograde methods. Analyses such as these highlight how the high degree of multiplexing (within a single sample) inherent in axonal BARseq enables identification of potentially interesting subpopulations.

### Axonal BARseq can identify cell types based on projection trajectory

Following previous analyses of auditory cortex (Chen et al., 2019) and other cortical structures (Harris and Shepherd, 2015), we manually clustered barcoded neurons into major cell types (Fig. 3A, SupFig. 6A-B). The top-level partition, between corticofugal (CF) and intratelencephalic (IT) classes, was based on the presence of subcortical projections descending below striatum, including the ipsilateral thalamus and the superior colliculus. IT cells were further divided into ITi and ITc, based on whether they had projections to the contralateral cortex. Barcoded somata were distributed across laminae and particularly enriched in mid layers (SupFig. 6C). Consistent with previous studies, CF somata were found predominantly in layer 5 (L5) and layer 6 (L6), whereas ITi and ITc somata were distributed across layers (Fig. 3B-C, SupFig. 6D, Chen et al., 2019; Harris et al., 2019; Winnubst et al., 2019; Muñoz-Castañeda et al., 2021). Thus, the projection patterns observed with axonal BARseq recapitulate those observed with conventional methods and with previous studies using BARseq.

CF neurons are divided into two major types (Harris and Shepherd, 2015): extratelencephalic (ET, also known as pyramidal tract/PT neurons) and corticothalamic (CT). ET and CT neurons are distinct in the laminar positions of their somata (Harris and Shepherd, 2015), axonal trajectory (Oh et al., 2014), gene expression (Tasic et al., 2016, 2018), and projection targets. ET neurons from the auditory cortex project to both the tectum (Chen et al., 2019) and higher order thalamic nuclei, including lateral posterior nucleus (LP), posterior limiting nucleus (POL), posterior intralaminar thalamic nucleus (PIL) and peripeduncular nucleus (PP; Cai et al., 2019; Harris et al., 2019). In contrast, CT neurons do not project to the tectum and mainly project to medial geniculate body (MG) in the thalamus (Llano and Sherman, 2008; Harris et al., 2019). In previous work (Chen et al., 2019) we distinguished ET neurons from CT neurons by the fact that only ET neurons project to the tectum. In the current experiment, however, we did not sample the entire tectum, and thus could not distinguish these two populations of neurons by the presence or absence of tectal projections. Instead, we exploited the high spatial resolution of axonal BARseq to partition neurons based on axonal trajectory.

Fig. 3D shows representative axonal trajectories to the thalamus of CF neurons. One group follows a dorsal route and travels through the reticular nucleus, whereas the second follows a ventral route. These two routes are consistent with the two axonal trajectories of CT and ET neurons, respectively (SupFig. 7A-D; Mitrofanis and Baker, 1993; Oh et al., 2014). We further combined axonal BARseq with immunohistochemistry to distinguish projections to different thalamic nuclei (SupFig. 7F-G). Consistent with the hypothesis that these two trajectories distinguish CT and ET neurons, neurons taking the dorsal route are concentrated in L5 (peak around 60% depth; Fig. 3B-C) and project to LP, POL, PIL/PP (Fig. 3D, SupFig. 7F-G). By contrast, neurons taking the ventral route are concentrated in L6 (peak around 90% depth; Fig. 3B-C) and project to the MG (Fig. 3D, SupFig. 7F-G). These results indicate that axonal BARseq can distinguish populations of projection neurons based on axonal trajectory.

### Diversity of laminar terminations across areas

The cortex is organized hierarchically, with primary sensory areas like auditory cortex representing the lowest level of the hierarchy. Feedforward projections–those that go up the hierarchy–often terminate in layer 4 of cortex, whereas feedback projections go to lower layers. Recently, Harris et al. (2019) reported that hierarchical position alone does not explain all of the connections made by a given area, and suggested that more complex models would be needed, with labels richer than just feedforward and feedback. However, the results of Harris and colleagues were based primarily on labeling of populations of neurons, leaving open the possibility that feedforward and feedback projections could be well-defined at the level of single neurons. We therefore studied the characteristics of areal and laminar projections across a large population of single neurons.

For the purposes of this analysis, we divided the cortex into four target areas: LatI, MedI, MedC and LatC (ipsilateral-lateral, ipsilateral-medial, contralateral-medial, contralateral-lateral, Fig. 4A). LatI/C mainly consists of auditory areas, whereas MedI/C mainly consists of higher-order visual areas (also called parietal associative areas; Lyamzin and Benucci, 2019; Wang et al., 2020). The majority of IT neurons from primary auditory cortex projected only to lateral (82.7%) areas, while almost no neurons (0.26%) projected to medial areas alone, and 17.1% projected to both. Thus, almost every neuron that targeted medial neurons also targeted lateral areas, but not vice versa, suggesting that information from neurons in the primary auditory cortex is obligatorily conveyed to primary and higher auditory areas (Lat), but only optionally to non-auditory areas (Med).

We next assessed laminar heterogeneity of projections to different cortical areas. The bulk-level laminar termination patterns of corticocortical projections are consistent with those observed in the Allen Mouse Brain Connectivity Atlas (Oh et al., 2014): projections to medial areas were largely localized to superficial layers (layer 1 to upper layer 2/3), whereas those to lateral areas were distributed across layers (Fig. 4B, SupFig. 8A-C). Because projection density in medial areas decreased strongly within layer 2/3 in our data set, as well as in data from the Allen atlas, we defined the border between “superficial” and “inferior” projections at the mid-point of layer 2/3 (Fig. 4C, SupFig. 8D-E). Note that this border, defined by axonal projection density, does not correspond to the conventional boundaries used to demarcate the borders between layers 1-6 defined by somatic properties.

There are several ways in which the bulk-level projection pattern (superficial targets for medial areas and mixed superficial and inferior targets in lateral areas) could arise. At one extreme, the population could be homogeneous: every neuron could project to every layer in every area. At the other extreme, the population could consist of highly specialized neurons, with each neuron projecting to only one layer and only one area. Between these two extremes, there could be subpopulations with specific patterns of projections. To disambiguate these possibilities, we examined the laminar terminations between medial and lateral areas at single-cell resolution. The neurons targeting medial areas projected mainly to superficial layers (as expected from the bulk data) and formed a relatively homogeneous population (Fig. 4D, SupFig. 8F). By contrast, single-neuron projection patterns in lateral areas were heterogenous, consisting of several diverse subpopulations.

In Fig. 4D, the diverse projection patterns are clustered for visualization purposes. Some neurons mainly project to superficial layers, representing a feedforward pattern; some mainly target deep inferior layers (Fig. 4D, *dot-labeled*), suggesting a feedback pattern. Additionally, there are neurons that project more diffusely across various layers, constituting an apparent mix of feedforward and feedback projections. An example neuron with projections mainly to superficial layers in MedI, but with diffuse projections to both superficial and inferior layers in LatI and LatC, is shown in Fig. 4E. Thus, it appears that the observed laminar projection patterns fall between the two extremes, with subpopulations showing specific patterns of projections. Our observations reinforce, at the single neuron level, the conclusions of Harris et al. (2019), who found a mix of feedforward and feedback projections in Cre-defined neuronal populations. These results suggest a need for more nuanced categories beyond the simple “feedforward” and “feedback” dichotomy.

Additionally we asked about the consistency of a laminar projection pattern at the single neuron level across cortical areas, using the superficial pattern as a representative example for this analysis. For neurons that project solely to a lateral target, the probability of projecting predominantly to the inferior layers (83.5%) is much higher than the probability of projecting to superficial layers (16.5%) (Fig. 4F, SupFig. 8G-H). However, for neurons that also project to a medial target, those same lateral projection probabilities nearly equalize (57.2% vs. 42.8%, respectively). Thus, the subpopulation of neurons that project to both medial and lateral areas are predisposed, but not determined, to project to superficial targets in both areas.

### Diversity of projections within an area

We next examined the relationship between soma laminar position and the laminar structure of axonal projections to different cortical targets. We divided Lat-projecting cells into three groups based on soma layer (Fig. 5A): upper (layer 1 to layer 2/3, in red), middle (layer 4 to upper layer 5, in orange), and deep (lower layer 5 to layer 6, in blue). Using these divisions, we found that at the bulk level, the deep (*blue*) somata mainly projected to deep layers (lower layer 5 to layer 6 as defined previously), whereas the upper and middle somata preferentially projected to upper-middle layers of their targets (Fig. 5B, SupFig. 9A-B). Notably, the upper border of deep projections is located around mid-layer 5 (Fig. 4D and 5E, Egger et al., 2020), which does not conform to the laminar boundaries.

We then compared the organization of these projections to LatI/C at the single-cell level. Three example neurons are illustrated in Fig. 5C-D. In these examples, and across the population (Fig 5E), projections from upper and middle somata to both areas (LatI, LatC) were largely enriched in upper-middle layers. The tendency of deep somata to project to deep targets is also reflected in Fig. 4D: 77.7% of deep-projection clusters (also see SupFig. 9C, *dot-labeled*) arose from deep somata. Moreover, we observed a marked correlation between the depth of a deep somata and the proportion of deep-projections (SupFig. 9D). Thus, the most superficial of the “deep” somata had more upper/middle-layer projections, and the proportion of upper/middle-layer projection gradually decreased with somatic depth (Chen et al., 2019). The correlation between soma depth and projection target was further validated by a *post hoc* analysis of injection variability across wild-type mice in the Allen Mouse Brain Connectivity Atlas (SupFig. 10). Also, the subset of callosal projections that terminate around the boundary between layer 4 and 5, and layer 1 in Fig. 4D, largely originated from middle-layer somata (84.8%, SupFig. 9C, *circle-labelled*). Similar projection patterns were observed from the middle somata (layer 4 and 5 IT Cre lines; SupFig. 9E-F), underscoring the consistency of laminar projection patterns between traditional Cre-based bulk tracing and axonal BARseq. Furthermore, we found that upper somata were less likely to project to ipsilateral striatum, and middle somata were more likely to project to Med cortex compared to the other two groups (Fig 5F-H, SupFig. 9G-I; see also Harris et al., 2019; Muñoz-Castañeda et al., 2021).

We next analyzed the fanout pattern of axonal projections within a target area. In LatC, upper somata tended to focus their axonal terminations in a small area (Fig. 5C-D, *red neuron*), whereas deep somata tended to project more diffusely (Fig. 5C-D, *blue neuron*), with middle somata exhibiting an intermediate pattern (e.g. two foci; see Fig. 5C-D, *orange neuron*). These observations are summarized across the population in Fig. 5I (*see also* SupFig. 9J-N, SupFig. 11). Taken together, axonal BARseq resolved systematic differences in the projections of layer-defined subpopulations of IT neurons in the laminar patterns of axon termination, their projection targets, and the projection patterns within a target.

## Discussion

We have described axonal BARseq, a highly multiplexed method for mapping neuronal projections with single-cell resolution. A key advantage of axonal BARseq over conventional optical methods is the large number of projections that can be mapped in a single brain. As a proof-of-principle, we used axonal BARseq to map the projections of more than 8,000 neurons from primary auditory cortex of a single mouse. Axonal BARseq represents an advance in spatial resolution over first-generation BARseq, which relied on bulk sequencing to read out projection barcodes. We used this large data set to systematically quantify the heterogeneity of auditory cortical projections to multiple targets. Additionally, we showed that axonal BARseq can be combined with routine immunohistology (SupFig. 7G).

The central challenge in multiplexed mapping of axonal projections is that the axons are densely packed and tangled together. When the distance between two axons approaches or falls below the limit of optical microscopy, the fidelity with which they can be distinguished using classical methods decreases. The greater the number of labeled axons, the greater the probability that two axons will be indistinguishable, and thus the greater the probability of error. Tracing errors are catastrophic because the error implies that an axon will be misattributed to the incorrect soma of origin. These challenges are particularly acute when tracing axons over long distances, because axons often travel in bundles. Such considerations limit the number of labeled axons that can be optically reconstructed within a single specimen.

Axonal BARseq circumvents these challenges by eliminating the need to trace axons. Instead, barcodes provide a direct means of associating the axon with its parent soma. Errors in appropriate attribution of a barcode (e.g. due to sequencing errors) are rare and, importantly, are not catastrophic because they are limited to a single rolony. Moreover, projections to distant targets can be assessed even without the need to trace the entire axonal path from soma to target. This enables efficient mapping of projections to multiple target areas, even if the targets are widely separated in space. The use of Sindbis virus for neural barcoding has been validated across various studies employing diverse methodologies (Kebschull et al., 2016a; Han et al., 2018; Chen et al., 2019; Gergues et al., 2020; Huang et al., 2020; Mathis et al., 2021; Muñoz-Castañeda et al., 2021; Hausmann et al., 2022). In particular, barcode transport is uniform even for long axons, as initially demonstrated in the locus coeruleus (Kebschull et al., 2016a), For a more complete discussion of the possible confounds due to viral toxicity, co-infections, degenerate barcode libraries, and other potential sources of error, please refer to Kebschull et al. (2016a) and Chen et al. (2019).

The high throughput of axonal BARseq is useful for at least four reasons. First, high throughput allows for statistical analyses using large numbers of single neurons, which has the potential to reveal statistical structure that is not evident with smaller sample sizes. Second, the fact that the samples come from a single animal is useful when individual animals are rare or valuable, such as for nonhuman primates, non-canonical model systems, transgenic animals, or animals that have been subjected to specific treatments or manipulations. Third, axonal BARseq allows for dense mapping of projections within a single brain, obviating the need to register all results to a single reference atlas. Avoiding registration eliminates the errors that arise from comparing across brains. Moreover, registration implicitly assumes that all brains are the same, whereas in some cases idiosyncratic differences between brains may be important. Finally, axonal BARseq can be combined with *in situ* gene sequencing (BARseq2, Sun et al., 2021), allowing high-throughput correlation of gene expression with projection pattern. These advantages make axonal BARseq uniquely useful for certain applications, such as studying the relative topography of projections.

Here, we used axonal BARseq to simultaneously trace different cell types within a single wild-type animal. By tracing the subcortical projecting cell types, ET and CT, we directly observed the differences between them, including the laminar distributions of their somata and their projection patterns in the thalamus (Fig. 3D, SupFig. 7F-G). We identified hundreds of IT cells projecting to multiple cortical targets, and were able to quantify the extent to which single neuron projections to different brain areas terminated in different laminae (Fig. 4D). We also found that IT cells can have focal or sparse patterns in their contralateral projections, with sparse projections originating from deep-layer cells and targeting deep layers (Fig. 5D and I). Focal projections originated from upper-middle layer cells with a different laminar distribution. Our results demonstrate that axonal BARseq can recapitulates previously observed differences between cell types and is highly effective in making novel discoveries in heterogeneous cell populations.

## Limitations and future developments

Axonal BARseq has several limitations compared with other methods. First, like conventional GFP-based tracing approaches, axonal BARseq reveals only axonal projections but not synaptic connections. To achieve synaptic resolution requires electron microscopy or visualizing synaptic markers using super-resolution microscopy or expansion microscopy. Second, the reconstructed axons may not have high resolution, as the axonal rolonies can be spaced as far as tens of microns apart. This means that branch points or even entire branches may be missed, which can affect the accuracy of the reconstructed neuronal projections. Although the neuroanatomical literature has traditionally placed a high premium on reconstructing the finest processes with high fidelity, for many applications the increased throughput—thousands of neurons per sample–may represent a useful tradeoff. For example, if the main interest is in the laminar distribution of axonal innervation (Fig. 4D and Fig. 5E), the fact that not all fine axonal processes are recovered may represent an acceptable compromise. It may also be possible to increase the density of axonal rolonies and thus the fidelity of reconstruction by improving the delivery of barcodes to axons (e.g. with a better carrier protein) and by refining the protocols for rolony recovery. Alternatively and additionally, it might be possible to combine axonal BARseq with either classic fluorophore-based tracing or brainbow (Livet et al., 2007). Finally, in the current work we did not attempt to resolve local axons near the injection site because of limitations of the current algorithms for automated base-calling of rolonies. However, newer algorithms may make it possible to resolve rolonies at high density.

There are several potential avenues for improving upon the current axonal BARseq method. First, axonal BARseq could be combined with conventional GFP-based tracing techniques. By combining the higher resolution of conventional single neuron tracing–the ability to resolve even the finest axonal branches– with the higher throughput of axonal BARseq. Second, axonal BARseq can be combined with the expression of endogenous genes, which would enable us to correlate projection patterns with transcriptomically defined cell types, allowing a better understanding of the differences in projection patterns both among and within cell types. Finally, we expect that it will be possible to increase the number of cells that can be analyzed using axonal BARseq. In general, the number of cells recovered by BARseq is determined by the size of the injection. In this study we restricted our injection to a single site, labeling a relatively small number of neurons. However, in previous work (Huang et al., 2020) we have barcoded more than 100,000 neurons in a single brain, and there is no technical barrier to labeling comparable numbers of neurons for axonal BARseq in future studies. Furthermore, Sindbis virus can infect diverse species including primates (Zeisler et al., 2023), so axonal BARseq could potentially be modified to map projections in many model systems, especially those in which conventional tracing-based approaches are impractical. Axonal BARseq thus has the potential to emerge as a powerful tool for massively multiplexed mapping of single neuron projections in diverse model systems.

## Methods

### Sindbis virus barcode library

The sindbis virus (SINV) barcode library used in this study was generated by the MAPseq core facility at Cold Spring Harbor Laboratory. The VAMP2nλ vector was derived from the MAPP-nλ SINV vector (Addgene #79785, Kebschull et al., 2016a) by replacing MAPP-nλ with VAMP2nλ. The VAMP2nλ consists of an nλ RNA binding domain, a V5-tag, and a mouse VAMP2 sequence. Oligos containing 30 nucleotide barcodes, including 28 random nucleotides and 2 fixed nucleotides, were synthesized by IDT. The barcoded virus library was produced as previously described (Kebschull et al., 2016b). Briefly, the digested oligos were inserted into SINV genomic vectors, transformed by electroporation for plasmid production. The plasmids were then linearized, and RNAs were *in vitro* transcribed using the mMESSAGE SP6 kit. The SINV genomic and package RNAs (Addgene #72309) were transfected into BHK cells using Lipofectamine 2000. After two days of expression, the SINV supernatant was filtered and ultracentrifuged for virus purification. The virus pellets were resuspended in 1xPBS and stored at −80℃. The VAMP2nλ SINV library exhibits a diversity of approximately 4 million barcodes, as determined by Illumina sequencing.

### Animal processing and tissue preparation

All animal procedures were approved by the Institutional Animal Care and Use Committee at Cold Spring Harbor Laboratory (protocol 19-16-10-07-03-00-4). Experiments were performed on 7-10 week-old male C57BL/6 mice (Charles River). The VAMP2nλ SINV library was injected into the primary auditory cortex using the NanojectIII (Drummond) at the following coordinates: −2.5 mm AP, ± 4.2 mm ML, 0.9, 0.6, 0.3 mm depth, with a volume of 150 nL per depth. The mouse used for axonal BARseq was injected at −4.2 mm ML. After 2 days of expression, animals were anesthetized and perfused with 4% PFA in 1X PBS. The samples were post-fixed at 4°C for a day and then transferred to sucrose gradients (10-15%, 20-22%, 30% sucrose in 1X PBS at 4°C) and frozen in OCT. The brains were cryosectioned at 20 µm thickness, mounted onto glass slides using UV-solidified glue (Norland Optical Adhesive NOA81, 8-10 s UV) to minimize section distortion or detachment during high temperature and chemical treatments.

### Rolony preparation

Before starting sample preparation, the sections were thawed and a hybridization chamber was placed on top. In the axonal rolony experiments, one section per chamber was utilized, while in the axonal BARseq experiment, two adjacent sections were used per chamber. To eliminate any residual fluids, chambers and samples were rinsed with water or reaction buffer before crucial reactions. For extended reactions or overnight reactions, humidified chambers were employed to prevent section dehydration. The catalog numbers and oligos utilized are listed in SupTable 2, respectively. The differences between this protocol and the BARseq protocol (Chen et al., 2019) are listed in SupTable 3.

#### Sample pretreatment

Samples were washed twice with 1% PBSTE (1XPBS with 1% Tween-20 and 5 mM EDTA) and incubated in 1% PBSTE at 65 °C for 8-9 min. Next, they were placed on ice for 2 min and washed twice with 1% PBSTE. The samples were then dehydrated in 50%, 70%, and 85% ethanol and incubated in 100% ethanol overnight at 4 °C. After two washes with 100% ethanol, the samples were washed twice with water and 1% PBSTE to smooth the chamber. They were briefly washed in 4 mM HCl to adjust the pH for pepsin digestion and then digested with 0.1-0.2% pepsin (w/v) in 4 mM HCl with 1 µM XC1215 at pH ∼3 at 33 ℃ for 30-40 min. It is important to note that the activity of the pepsin solution varies batch-to-batch and the activity of each batch was tested. Similarly, the pH of the solution was monitored as a low pH results in high nuclear background during *in situ* sequencing, while a more neutral pH leads to low pepsin activity. Finally, the duration of pepsin digestion was closely monitored, as over-digestion can cause tissue/rolony degradation/tearing or falling off, while insufficient digestion can lead to low permeability and low rolony density.

#### Reverse transcription

After digestion, samples were washed twice with 1% PBST and then washed in 1X SSIV (SuperScript IV) buffer containing 0.4 µg/µL BSA and 5 mM DTT for 5-15 min at room temperature. Reverse transcription was performed on the samples using 1 µM XC1215, 20 U/µL SSIV, 500 µM dNTP, 0.2 µg/µL BSA, 1 U/µL RiboLock RNase Inhibitor, and 5 mM DTT in 1X SSIV buffer at 45 ℃ for approximately 4 hours. Samples were then transferred to a new reaction mix and incubated overnight at 45 ℃.

For the reagent comparison experiment (SupFig. 1B-C), samples treated with RA (RevertAid H minus reverse transcriptase) were washed in 1X RA buffers containing 0.4 µg/µL BSA for 5-15 min at room temperature. Reverse transcription was performed on the samples using 1 µM XC1125, 20 U/µL RA, 500 µM dNTP, 0.2 µg/µL BSA, and 1 U/µL RiboLock RNase Inhibitor in 1X RA buffer at 37 ℃ for approximately 4 hours. The samples were then transferred to a new RA reaction mix and incubated overnight at 37 ℃.

After reverse transcription, the samples were washed with 1X PBS and crosslinked with 25 mM BS(PEG)9 in 0.2% PBST for 30 min at room temperature. They were then washed with 0.2% PBST (0.2% Tween) and incubated in 2 mM lysine in 1X PBS for 30 min.

#### Gapfilling

After crosslinking, the samples were washed with 0.2% PBST twice and with water twice, and then gapfilled with 100 nM padlock probe LYO5, 0.5 U/µL Ampligase, 50 µM dNTP, 0.4 U/µL RNaseH, 50 mM KCl, 20% formamide, and 12.5 mU/µL Phusion in 1X Ampligase buffer at 37 ℃ for 30-40 min, and 45 ℃ for 45 min. To prevent Phusion from reacting with the padlock, the reaction mix was kept cold and Phusion was added immediately before the reaction. In the reagent comparison (SupFig. 1B and D), padlock probe LYO5 was replaced by XC1164.

#### Rolling circle amplification (RCA)

After gapfilling, the samples were washed thoroughly with 0.2% PBST and rinsed with water. They were then incubated with RCA mix (1 U/µl EquiPhi29 polymerase, 0.25 mM dNTP, 120 µM aadUTP, 0.2 µg/µL BSA, and 1 mM DTT in 1X EquiPhi29 buffer) at 37 °C overnight. After incubation, the samples were washed with PBS once and crosslinked with 25 mM BS(PEG)9 in 0.2% PBST for 15 min at room temperature. They were then washed with 0.2% PBST twice and quenched with 1M Tris pH 8.0 for 30 min. This RCA-crosslinking step was repeated two more times. After three rounds of RCA, the samples were crosslinked with 25 mM BS(PEG)9 in 0.2% PBST for 30 min at room temperature. They were then washed with 0.2% PBST twice and quenched with 1M Tris pH 8.0 for 30 min.

### Axonal barcode detection comparison

To measure the sensitivity of rolony preparation, we compared it to FISH, a standard method with high single-molecule sensitivity. In these experiments, rolonies were hybridized with fluorescence-conjugated probes. After rolony preparation, the samples were hybridized with 0.25 µM probe XC92 in 2X SSC, 10% formamide for 15-30 min at room temperature. Any excess probes were washed away three times with 2X SSC, 10% formamide, and three times with 0.2% PBST.

In the experiments used to compare rolony preparation and FISH, the FISH samples were pretreated in the same way as the rolony preparation samples. After digestion, they were washed, and FISH was performed using GFP probes and the RNAscope kit according to the manufacturer’s protocol.

Quantification was performed using max-projected and stitched images. Similar regions of interest were manually selected in the AudI/AudC/Thal/VisI for each brain section, avoiding somatic rolonies. Rolony counts were measured using ‘Find Maxima’ with fixed prominence in Fiji, and density was calculated by dividing by the area size. Rolony densities were normalized to the density of the same region in neighboring SSIV + LYO5 samples. The median of the normalized density was calculated from 2-4 regions per section.

Interestingly, we found that a 2 nt length difference between padlock probes LYO5 and XC1164 significantly affected rolony signals. This may be because template length affects Phi29 efficiency during RCA (Joffroy et al., 2018). While the modified protocol generated more axonal rolonies, it was less cost-effective for producing somatic rolonies. Therefore, for somatic barcode sequencing, the original BaristaSeq protocol (Chen et al., 2018) is sufficient due to the high abundance of somatic barcodes.

### Axonal and somatic rolony comparison

Probe-hybridized samples in SupFig. 1E-F were used for comparison of axonal and somatic rolonies. For each brain section, 10-20 somatic areas were manually selected in AudI, and somatic rolony intensities were represented by the maximum intensity of each somatic area. In the same stitched images, 6-7 300×300 pixel ROIs were manually selected for axonal rolonies, avoiding somatic rolonies. Within each ROI, axonal rolonies were identified using ‘Find Maxima’ in Fiji, and axonal rolony intensities were represented by the intensity of the maxima. The median intensities of axonal and somatic rolonies were calculated for each section, with background subtraction.

### In situ sequencing

Axonal BARseq samples were split into seven rounds of rolony preparation (SupTable 1). For each round, the samples were divided into two batches for *in situ* sequencing. After rolony preparation, the samples were incubated in 2X SSC, 80% formamide at 65 ℃ for 15 min. 2.5 µM sequencing primer LYO23 was hybridized to the rolonies in 2X SSC, 10% formamide for 15-30 min at room temperature. Any excess primers were washed away three times with 2X SSC, 10% formamide, and three times with 0.2% PBST.

*In situ* sequencing was performed using the HiSeq Rapid SBS Kit v2 (Illumina). The reagents used in this process included the Universal Sequencing Buffer (USB), Cleavage Reagent Mix (CRM), Cleavage Wash Mix (CWM), Incorporation Master Mix (IMT), and Universal Scan Mix (USM). Before the first cycle, the samples were washed with USB at 60 ℃ for 4-5 min twice. Then, they were incubated with CRM at 60 ℃ for 5 min. The samples were washed three times with CWM, 1% TT (20 mM Tris pH 8.0, 1% Tween-20), and twice with PBS. Next, they were blocked with iodoacetamide (9.3 mg tablet in 2 mL 1XPBS) at 60 ℃ for 4-5 min and washed with 0.2% PBST three times. For each sequencing cycle, the samples were washed with USB at room temperature twice and incubated with IMT at 60 ℃ for 4 min. They were then washed with 1% TT with 5 mM EDTA once, and 1% TT at 60 ℃ for 4 min 3-5 times. The samples were incubated in USM and were ready for imaging. After imaging, the samples were washed with 1% TT three times and USB twice, incubated with CRM at 60 ℃ for 4 min, and washed with CWM. In the later sequencing cycles, the C-channel often had a high level of nonspecific background; additional 1% TT washes were included to decrease this background. In round 1 of this dataset, the samples did not receive CRM treatment before the iodoacetamide incubation prior to Seq01. Additionally, an extra iodoacetamide treatment was applied after the first CRM step following Seq01 imaging.

### Immunohistochemistry

After the final sequencing cycle (Seq17), the samples were treated with CRM and CWM to remove any remaining sequencing signals. They were then blocked with 5% BSA in 1X PBS and incubated with a vGlut2 antibody (1:500) in 2% BSA in 1X PBS at 4 ℃ for 2 days. Following washes with 0.2% PBST, the samples were incubated with a secondary antibody (1:1,000) in 2% BSA for 2-4 hours at room temperature. After additional washing, the samples were stained with DAPI and imaged using USM.

### Microscopy

Images were obtained using a Nikon TE2000-E microscope equipped with an X-Light V2 spinning disk (Crest Optics), Prime 95B camera (Teledyne Photometrics), and LDI-7 laser diode illuminator (89North). A 20X Plan Apo objective (Nikon) was used for all experiments. It is important to note that factors such as optical distortion and uneven illumination in the microscope system can affect the sensitivity and accuracy of axonal BARseq. Images were acquired using µManager 1.4 (Edelstein et al., 2010). All images were taken as z-stacks with the following settings: 0.55 µm per pixel, 12-bit depth, a total of 17 stacks with 3 µm intervals, and 15% overlap for tiling. The lasers and filters used for each channel are listed in SupTable 2. Briefly, each nucleotide was imaged in a separate channel during sequencing. We found that maximum intensity projection preserved most of the signal while resulting in smaller file sizes and reduced computational demands during analysis. As a result, we converted the z-stacks to max projections.

### Image processing and rolony identification

The general workflow for data acquisition and analysis is described in SupFig. 2A. In order to strike a balance between data output and imaging time, we opted to image selected target areas of auditory cortex. The target areas were initially identified through manual selection and subsequently registered to the corresponding coronal section obtained from the immunohistochemistry step to ensure accurate positioning. Due to the selection of target areas, processes outside these regions were not included in the current study.

The image processing workflow for *in situ* sequencing is described in SupFig. 2B. To reduce fixed pattern noise during *in situ* sequencing, we subtracted the 3rd lowest intensity plane of the z-stack from the maximum projection image. Local background subtraction was performed by taking advantage of the fact that pixels without a barcode have discontinuous intensity profile along the z-axis. This process effectively removed local signal distortions and backgrounds such as uneven illumination, nuclear, and tissue background (SupFig. 2C-D). However, while this method was effective for axonal rolonies in target areas, errors were encountered in pixels around the somata due to the high signal density and the aberrant intensity distribution along the z-axis compared to single rolonies. To correct for bleed-through, uneven channel intensity, and intensity decay across sequencing cycles, we based intensity corrections on local maxima for each experiment. To decrease variability between individual batches, we used z-scores for intensity correction. Rolonies were typically between 3-7 pixels in diameter on the maximum projection images. Therefore, we identified local maxima within a 5-pixel diameter range for each sequencing cycle. A local maximum was considered a rolony location if it met the following criteria: (1) in the z-stack, the slides with the highest intensity were neighboring slides (e.g. the 1st max intensity slide was next to the 2nd/3rd max intensity slides); (2) the channel with the local maximum had the highest intensity before and after image corrections; (3) the channel intensity passed the intensity ratio filter (i.e. 2nd max/1st max < 0.95); (4) the max channel intensity passed a threshold. To improve the accuracy of matching rolonies during base-calling, we calculated the subpixel locations of local maxima using interpolation.

For immunohistochemical experiments, maximum projections of image tiles were stitched into whole coronal sections using phase correlation, with max projection in the overlapping region.

#### Tile alignment and stitching

The workflow for alignment and stitching is described in SupFig. 3A. The alignment process consists of two steps: (1) pre-alignment using stitched images; (2) point cloud registration for individual tiles.

During pre-alignment, tiles from the same image were initially stitched using imaging positions and then aligned across sequencing cycles using phase correlation. We used imaging position-based stitching to avoid errors from intensity-based algorithms. Additionally, stitched images were aligned to 1-3 sequencing cycles to minimize errors. After pre-alignment, rolony coordinates were aligned to the reference or neighboring sequencing cycles using a projective/affine transformation (SupFig. 3B). The transformation matrix between point clouds was calculated using a frequency-based algorithm. To reduce the impact of tissue distortion during sequencing, we used mid-sequencing cycles (Seq08/09) as reference cycles. Vis of section #76 was excluded during alignment.

For stitching, we combined and aligned the rolony coordinates from nearby tiles across sequencing cycles to the neighboring tiles. To minimize errors, we stitched tiles with a lower number of rolonies to tiles with a higher number of rolonies.

#### Rolony base-calling

Our rolony base-calling pipeline allowed for a degree of error during alignment. No non-linear transformations were performed during alignment. Base-calling was performed by matching nearby local maxima (dots) across sequencing cycles (SupFig. 3C). Dots were first given unique IDs in each sequencing cycle, and dots from later cycles were one-to-one matched to the closest available dots in the previous cycle within a 5-pixel range. The sequence of dots was then assembled, and the nucleotides associated with the dots were identified as the rolony barcode.

During sequencing, rolony signals may be lost, shifted, or near a strong nonspecific noise signal. To maximize continuity in the sequencing results, each rolony was matched to rolonies in three previous cycles. To assemble the sequence, matches were merged sequentially from a 0 to 3 cycle interval in ascending order (e.g., 3-4, 2-4, 1-4, 4-5, 3-5, 2-5, etc.). Non-base-called nucleotides were assigned to intervals when two matching cycles were not consecutive. If there was a disagreement between the current match and the existing sequence, the previous sequence was duplicated to include the new match (SupFig. 3C, blue). Barcodes with more than three continuous non-base-called nucleotides were discarded. However, due to the high density of signals and resulting higher error rate near the injection site, we excluded rolonies close to the soma (within 20 µm of ≥ 35 soma pixels in > 2 cycles) from the analysis.

#### Soma base-calling

Since barcoded somata were larger in size compared to individual rolonies, we base-called somata using pixel location rather than local maxima. We identified the barcodes as the channels with the highest intensity across sequencing cycles using stitched images from the AudI. For technical reasons, the stitched images used for soma base-calling only included one tile for overlap areas.

During later sequencing cycles, we noticed that barcoded somata showed phasing signals, but single rolonies did not. This may be because somata contain a larger number of barcoded single-stranded DNA, and the protocol was not optimized for soma barcode sequencing. To digitally correct this, we subtracted the pixel intensity from the previous cycle (50% for maximum intensity and 100% for the rest), which improved the signal-to-noise ratio (SupFig. 3D).

#### Registering image to brain volume

We identified and imaged targeted brain regions separately for *in situ* sequencing. After alignment and stitching, we registered the images to the whole coronal section using nuclear signals by phase correlation. We then aligned the coronal sections into a 3D volume using control point pairs. These point pairs were selected manually between nearby sections, and displacement fields were generated from polynomial2 and piecewise linear transformation within manually defined limits and corrections. One brain section (section #108) was excluded due to severe distortion.

#### Codebook and lookup table

We used the results of the *in situ* sequencing to construct a list (codebook) of infected barcodes. The two sources of barcode combinations are axons and soma. As this study focused primarily on axonal projections, we chose only axonal barcodes for our codebook (SupFig. 4A).

We tolerated 1-2 nt errors in our rolony base-calling protocol, since the base-calling process could have errors either due to single mutations during sample preparation or misalignments during analysis. This ensured the accuracy of the codebook/lookup table and minimized data loss.

As described above, the rolony base-calling procedure can base-call a rolony multiple times, resulting in a set of barcodes with and without errors. To construct our final codebook for this dataset, we made the following assumptions: (1) a true barcode can be found in multiple rolonies (≥ 3 rolonies); (2) a true barcode has a higher count than its erroneous versions; (3) there are no pairs of true barcodes within 1 Hamming distance (<0.1%, SupFig. 4B); (4) a true barcode consists of different nucleotides (< 14 same nucleotides) and meets length limits (with ≥ 13 of 15 nucleotides base-called, ≤ 3 continuous non-base-called nucleotides). Based on these parameters, our codebook consisted of 13,919 barcodes.

Lookup tables were used to match individual axonal and somatic barcodes to the codebook. A Hamming distance of 2 was set as a cutoff to match axonal and somatic barcodes to the codebook barcodes. Barcodes that matched more than one codebook barcode within the minimum Hamming distance were discarded. During this process, non-base-called nucleotides were treated as a match to all four nucleotides if there was no mismatch; otherwise, they were treated as a mismatch. All nucleotides were included in the Hamming distance calculation at this step. To minimize misassignment, we constructed the codebook and lookup table before filtering, as eliminating a potential barcode early on may result in its axonal rolonies being assigned to another barcoded cell within the maximum Hamming distance.

#### Axonal barcode correction

During axonal base-calling, it was possible for a single axonal rolony in one sequencing cycle to link to more than one rolony in another sequencing cycle. This can result in (1) one rolony belonging to multiple different barcodes; (2) the same axonal rolony being called multiple times and linked to different rolonies in other cycles, but belonging to the same barcode. To address these issues, we took the following steps: (1) a rolony in a cycle linked to more than one barcode was excluded and the cycle was assigned as non-base-called; (2) barcodes that did not meet the requirements for length and interval were excluded; (3) barcodes with similar sets of rolonies were condensed into the one with the most base-called digits.

The Hamming distance between a pair of barcodes was calculated as the total number of mismatches between them. By default, non-base-called nucleotides were treated as a match to all four nucleotides. In this SINV library, the 9th and 10th nucleotides were fixed and, therefore, excluded from the Hamming distance calculation, unless stated otherwise.

#### Soma identification

Soma barcode counts were determined by counting the number of pixels associated with a specific barcode at the injection site (AudI). However, these counts alone were not always reliable for identifying barcoded somata in our current setup, potentially due to the following factors: (1) some somata were cut and split into two neighboring sections during sectioning; (2) loss of surface area of the section during sample preparation (e.g. due to over-digestion by pepsin); (3) weak signals in deep areas of the section due to insufficient permeabilization during sample preparation; (4) low soma barcode counts in some cells; (5) base-called areas appearing smaller than they should be due to alignment, stitching errors, and phasing; (6) soma base-calling being sensitive enough to identify dendritic, and occasionally axonal, rolonies in AudI.

To identify valid soma locations, we identified the brain section with the highest sum intensity of soma pixels as the soma section, and within this section, we identified the XY coordinates of the brightest pixels as soma locations. A soma needed at least 80 counts of its barcode within 100 µm of its location. Barcodes that did not meet these criteria were identified as barcodes without soma locations. In the registered data, the median distance between somata and the injection center is 267 µm.

#### Barcode Filters

##### Filtering out error-prone barcodes

To reduce the number of nonspecific barcodes, we applied the following filters: (1) barcodes with > 6 continuous identical nucleotides were excluded (152 out of 13,919); (2) to minimize sequencing errors arising from bleedthrough during imaging, barcodes with more than 14-nt in Ch1/2 or Ch3/4 were excluded due to the similarity of excitation wavelengths (127 out of 13,767); (3) barcodes with more rolonies with 1 or 2 mismatches compared to no mismatches were excluded (82 out of 13,640). Non-base-called nucleotides were treated as a match to all four nucleotides at this step. Only barcodes that passed the count filter (1) with ≥ 10 rolonies in at least one target region, (2) with ≤1,000 and ≥ 3 axonal barcode counts, and (3) with ≤ 7,000 somal barcode counts were included for analysis (9,185 out of 13,558 were included).

##### Secondary infection exclusion

We observed secondary infection in target brain regions, and most of the infected cells had a glial morphology. We manually identified 17 barcodes from these cells in all regions except AudI. Barcodes within a 4-Hamming distance of these identified barcodes were excluded from the analysis (92 out of 9,185).

##### Repeated rolony exclusion

To avoid double-counting, we excluded repeated rolonies in overlapping imaging fields, such as the cortex and striatum. Specifically, in these overlapping areas, we only included one copy per barcode from different fields (exclusion range < 25 µm).

##### Floating rolonies identification and exclusion

We observed that rolonies could sometimes float out of the soma and settle within a surrounding area. Among barcodes without a soma location, we also observed this floating rolony effect. Since the soma locations were unknown, we could not use the soma section to exclude these floating rolonies. Therefore, we used an alternative method to identify sections with floating rolonies. We used two criteria for identifying these sections: (1) the slide (and sometimes its neighboring slide) was the only one with rolonies in specific areas, and (2) the rolonies on the slides were widely and sparsely distributed. To identify rolonies that meet criterion (1), we excluded rolonies with neighbors (< 140 µm) in other sections (> 1 section away). To test whether criterion (2) was met, we identified a section to have enough rolonies that were far apart (≥ 3 rolonies/clusters with a distance beyond 50 µm). For barcodes with more than one such section, we selected the one with the widest rolony coverage. We used AudI, Thal, and Vis to find slides with floating rolonies.

We used this algorithm to identify sections with floating rolonies in cells with and without soma locations. Verification using cells with soma locations showed that the algorithm identified floating rolonies in 37.1% of cells. Within these positive barcodes, the algorithm had an accuracy of 94.8% for identifying the range of sections for the soma (± 1 section). For cells without soma locations, the algorithm detected floating rolonies in 14.9% of barcodes. We manually validated 75 positive barcodes and found a false positive rate of 22.7%. We achieved 100% accuracy for identifying the range of sections for floating rolonies. False positives would result in the exclusion of true rolonies in a 40-60 µm area in selective targets, but since projections usually extend more than 200 µm, this had a limited effect on downstream analysis.

To exclude floating rolonies, we excluded axonal rolonies in AudI, Thal, and Vis from 2-3 sections around the soma sections for barcodes with soma locations, and from the floating rolony section for barcodes without soma locations. After applying these exclusions, 8,838 barcodes passed the count filters.

##### Non-neural cell exclusion

Although the strain of SINV we used preferentially infects neurons versus non-neuronal cells, this preference is not complete, so some non-neuronal cells are also infected. In this study we did not distinguish between neuronal and non-neuronal cells using cell-specific markers. To exclude non-neural cells from our analysis, we applied distance and counts criteria. Specifically, cells needed to have ≥ 3 axonal rolonies ≥ 200 µm from the soma or center of axonal rolonies in AudI, and 50 cells were excluded using this criterion. This criterion was applied because non-neural cells typically do not have long projections. It is worth noting that this process may also filter out neurons with short local projections.

##### Additional filtering after CCF registration

After registering the data to the CCFv3 reference frames we applied the following additional filters. We first deleted rolonies outside the CCFv3 brain area and performed additional floating rolony elimination in the hippocampus, ventricle and fimbria of CCFv3. Next, we set a minimum rolony counts for five major targets: 5 for the ipsilateral/contralateral cortex and thalamus, 3 for the striatum and midbrain. After filtering, we excluded 167 barcoded cells with < 10 counts in any imaging region, as well as 16 non-neural cells. It is worth noting that these steps are optional and can be skipped. After the above-mentioned steps, we identified 8,620 barcodes, including 3,700 with soma location. Four barcodes had single-digit non-base-called nucleotides.

##### Manual assessment of base-calling results

To assess the sensitivity and accuracy of our automated base-calling pipeline, we compared it to manual base-calling. To evaluate the sensitivity of axonal rolony base-calling, we calculated the percentage of base-called rolonies in 17 randomly selected Seq14 images from target areas (3-6 ROIs per image, 300 x 300 pixels). Sensitivity was 44.5% ± 9%. To estimate the accuracy of axonal rolony base-calling, we randomly selected 18 barcodes and found that 0 out of 60 (0%) rolonies had > 2 nt disagreements between the codebook and evaluator.

To evaluate the efficiency of soma base-calling, we manually selected 50-112 somata per image in 7 randomly selected sequencing images, and found that 63.9% ± 7.6% of somata were base-called. To estimate the accuracy of soma base-calling, we randomly selected 40 barcoded somata and manually base-called them, and found that 5 (12.5%) had > 2 nt disagreements between the codebook and evaluator. To evaluate the accuracy of soma location, we randomly selected 90 barcoded somata and found that 79 (87.8%) were in agreement with the evaluator’s assessment. It is worth noting that there was high signal density near the injection site, which may have contributed to some uncertainty in the evaluator’s assessments.

#### Registering to Allen mouse brain CCFv3

To align the 3D data volume with the CCFv3 (Wang et al., 2020), we used a manual linear registration process. We then applied nonlinear adjustments to the coronal plates using control point pairs, similar to the method used for image registration to the brain volume. We used the Nissl reference map for this process. All reference maps used in this study (Nissl, average template, and annotation map) had a voxel resolution of 25 µm.

#### Evaluation of CCFv3-registration

To evaluate the accuracy of CCFv3-registration, we used a manual selection-based approach instead of intensity-based algorithms to minimize the impact from staining variabilities and uneven illumination. The edges of specific brain regions were manually selected on registered images and compared to their corresponding boundaries in CCFv3 (SupFig. 4D). To avoid bias, images and hemispheres were randomly chosen during the selection process. vGlut2 images (25 µm/voxel) were utilized to visualize the boundaries of brain regions. Six edges were chosen, including the outer edge of isocortex, outer edges of midbrain/thalamus/hypothalamus, outer edges of hippocampal formation (excluding entorhinal area), fasciculus retroflexus, mammillary related areas, and lateral edges of striatum/amygdala. The median shortest distance between the manually selected edge and the corresponding area boundary in CCFv3 was calculated.

The lateral edges of the registered striatum exhibited a lateral shift compared to CCFv3 and this discrepancy potentially led to errors in assigning striatal rolonies to lower layers of the cortex. To assess the impact of this issue, we manually annotated the ipsilateral striatum using vGlut2 registered images. Among the 6,878 LatI-projecting cells, a total of 454 cells had striatal rolonies assigned to LatI, with 5.01 ± 5.35% (mean ± SD) of the rolony per cell. This analysis suggests that the influence on the assessment of cortical rolonies distribution in this paper was minimal.

#### Cortical flatmap and ML/AP/depth-coordinates

To compare projections across cortical regions and hemispheres, we generated a lookup table for a cortical flatmap from the CCFv3 (SupFig. 4E-J). The concept of this flatmap is similar to that of (Wang et al., 2020), but with some differences. The flatmap coordinates consisted of three axes, with one axis oriented in the same direction as the cortical columns and the other two on a plate perpendicular to the columns. We defined the outer and inner cortical boundaries using the outer boundaries of layers 1 and 6, respectively. To determine the direction of the cortical columns, we calculated the lines from each outer boundary voxel to the closest inner boundary voxel. The depth percentage was the percentage of cortical depth along individual column lines. Due to cortical curvature, the distance between two column lines may vary at different depths (i.e., the distance is larger in upper layers compared to lower layers). We chose the mid-cortical plate (∼50% depth) as the reference plate for the other two axes. The values of the other two axes were calculated as the cumulative sum of voxel-to-voxel distances on the plate, and column lines were assigned to the values of their intersections at the reference plate. To obtain continuous, smooth values in all three axes, we applied an average filter.

Following general practice, we divided the reference plate into medial-lateral (ML) and anterior-posterior (AP) axes. Our goal in creating the flatmap was to simply flatten the cortex for physical distance, rather than attempting to represent biological gradients. We determined the definition of each axis with the following considerations: (1) rotating the brain around the x-axis in 3D space changes the direction and voxel value of the AP-axis; (2) similar to earth mapping, it is impossible to get a flat and continuous cortical plate without distorting the direction of the axes or the point-to-point distance due to cortical curvature. Although the midline is generally considered the ‘medial’ part of the brain, we found that setting the midline as a fixed value to flatten the cortex with this algorithm caused relatively more distortion near the lateral region. Therefore, we used principal component analysis (PCA) on the reference plate of the right hemisphere to define the 1st axis as AP and the 2nd axis as ML, and verified this visually. The contour line of the median AP and ML values was used as the reference line for flattening. The AP value was assigned as the distance to the AP reference line on the reference plate, and the ML value was assigned as the distance to the ML reference line with the minimum AP value change along the reference plate. The minimum value in both axes per hemisphere was set to 1. This flattening method was unable to differentiate between cortical regions folded towards the midline and the increased distortion at the lateral edge. However, since these were not the target areas of this dataset, the flatmap algorithm did not adjust for them. Additionally, this flatmap preserved the subtle voxel difference between the left and right hemispheres in CCFv3.

We used these lookup tables with interpolation to convert the registered rolony and soma locations into ML/AP/depth coordinates. To minimize interpolation error near the edges, we applied a 50 µm non-zero average filter to the outer edge of the cortex. For visualization purposes, we assigned the ML-values of the left cortex as negative and the right cortex as positive, and excluded the range of the AP-axis without cortical rolonies. During this process, two of the 3,700 somata were excluded, resulting in a total of 3,698 somata.

After registration, we defined the areas of four major projection targets (cortex, thalamus, striatum, and midbrain) using the CCFv3. We further divided the cortex into ipsilateral and contralateral cortices using the midline (SupFig. 6A). We drew the brain boundaries based on the parents of the 11th level of CCFv3. In the cortical flatmap, we represented the area boundaries by boundaries between 45-55% cortical depth, unless specified otherwise.

### Cortical layer gradience

In addition to utilizing cortical layers as anatomical markers, we computed gradients within each layer to detect potential biological variations. Cortical layer positions were determined using the Allen CCFv3 at a voxel resolution of 25 µm. The cortex data was then transformed into flatmap coordinates, and gradients within each layer were calculated for every ML-AP position along the cortical depth.

### Simulated retrograde tracing

We manually identified three injection centers for simulated retrograde tracing on the flatmap. We defined a range of 300 µm around the center as the injection/patch region. To be considered positive for retrograde tracing, a cell must meet the following criteria: (1) be from the IT class, (2) have ≥ 10 rolonies within the patch region, and (3) have a soma location. To identify cells that specifically project to a contralateral patch, the patch/CtxC rolony count ratio must be ≥ 75%.

### Single-cell tracing reconstruction

We created reconstructions by connecting the registered xyz-coordinates of rolonies and soma from the same barcoded cells. We first connected data points (including rolonies and soma) to their closest neighbor to form clusters, then connected each cluster to the nearest cluster via the closest data points until all clusters were connected. We set the maximum distance for connecting two data points at 1000 µm. We only included cells with soma location in the reconstruction, and dilated the somata for visualization purposes.

We computed the transparent outline of the brain using the CCFv3 annotation map. In coronal view images, we excluded stacks anterior or posterior to the current dataset (e.g., olfactory bulb and cerebellum) for visualization purposes.

### Grouping major cell types

We divided barcoded neurons into CF and IT cells based on projections to the ipsilateral thalamus and superior colliculus. CF cells had projections to either the Thal or SupCol (Fig. 3A) with minimum rolony counts as described above; all other cells were assigned as IT cells.

We used a two-step approach to classify Thal+ cells as either ET or CT cells. In the first step, we grouped cells based on the presence of rolonies in the striatal-thalamic fiber and thalamic reticular nucleus. The AUD axons in this region could be divided into two bundles, a dorsal and a ventral bundle in the coronal view and corresponding to axons from CT and ET cells (Mitrofanis and Baker, 1993). We manually defined a region of interest in this region using registered xyz-coordinates (x: 8,250-9,000 µm; y: 3,500-5,000 µm; z: 6,750-7,500 µm; red region in SupFig. 7C) and divided the rolonies within it into two groups based on their y-axis location: the top half were classified as CT cells and the bottom half were classified as ET cells. We then assigned each individual rolony to the most frequent group of its nearest 10 neighbors and followed this by assigning each barcoded cell to the most frequent group. This process was repeated until convergence or after 100 iterations. The cell type for each barcoded cell was represented by the most frequent group of rolonies within the region of interest. The results of this initial grouping are shown in SupFig. 7D (930 of 1,134 Thal+ cells were assigned to the CT/ET group; CT: 581; ET: 349). In the second step, we assigned all thalamic rolonies to the most frequent group of their nearest 10 neighbors, and the cell type for each barcoded cell was represented by the most frequent group of thalamic rolonies. 58 of the 930 cells were assigned to a different group in this step compared to the first step. The final results of the ET/CT grouping are shown in SupFig. 7E (CT: 713; ET: 421). Overall, this approach was able to classify the cell types for 91.2% (321/352) of Thal+ cells that project to the thalamus and the superior colliculus as ET cells and 8.8% (31) as CT cells, indicating that axonal BARseq can effectively identify cell types. We also observed that CT cells have rolonies in the striatum due to their axons traveling through the region to reach the thalamus. Visual examination showed that the majority of cells with rolonies in the striatum belong to the ET group (SupFig. 7F).

IT cells were divided into two subtypes: ITi and ITc. IT cells with ≥ 5 rolonies in the contralateral cortex were assigned to the ITc group, while the rest were assigned to the ITi group.

### Visualization and quantification of soma laminar distribution

To visualize the distribution of somata across groups, we plotted them using flatmap coordinates: x-axis, ML; y-axis, depth %. The plotting sequences were randomly shuffled across groups. Note that the ratio of the x-axis to the y-axis is not equal for visualization purposes.

To quantify the proportion of somata from different groups at different depths, we binned the soma depths into 5% bins and calculated the percentages for each bin: (group count) / (all group counts) × 100%.

### Visualization and quantification of rolony laminar distribution

To visualize the rolonies in the CCFv3, we plotted them in registered xyz-coordinates using a coronal view. The brain outline was shown in gray.

To visualize the laminar distribution in the cortex, we presented the data as heatmaps, unless otherwise specified. The frequency of rolony depth within a region was determined for each cell or bin, utilizing bin sizes corresponding to either 1% or 5% of the depth, or fractions of 1/2 or 1/4 of the layers. The darkness of the grid represented the relative probability.

### Cortical rolony analysis

In our experiments, we excluded cortical rolonies that were deeper than 95% due to their proximity to the fiber tract and potential registration errors. Most infected somata were localized to the middle layers. We excluded rolonies near the somata based on their xyz-coordinates, which may create uniform exclusion across cortical depth. Thus, for the cortical analysis (Fig. 4-5, SupFig. 8-11), we excluded rolonies that were located < 95 percentile of the injection center on the ML-AP plate (indicated by the black disk; the injection center is the median soma location). As a control for this exclusion, we also excluded rolonies in the same region on the other hemisphere (referred to as the LatC local-exclusion control).

Although the carrier protein VAMP2nλ was based on VAMP2, which localizes presynaptically, we found barcodes in dendrites as well. We excluded rolonies close to the somata using two filters as described above, so we believe the impact of dendritic barcodes on the cortical analysis is minimal.

The separation of Med and Lat targets was described in Fig. 4A, and we required a minimum of 5 rolonies per target for LatI/MedI/MedC/LatC-projection unless stated otherwise.

To calculate the normalized density of projections across cortical layers, we determined the thickness of each layer or bin. Bin thicknesses were computed using the cortical flatmap, excluding depths exceeding 95% as previously described. As the thickness of layers can vary across different cortical areas, we performed these calculations separately for the four cortical areas. The boundaries of each area were defined based on the locations of 95% of the rolonies within that area.

### Cortical quantification of auditory GFP bulk tracing

The registered GFP bulk tracing data was downloaded from the Allen Mouse Brain Connectivity Atlas (Oh et al., 2014), with a voxel resolution of 25 µm. The cortical data were first transformed into flatmap coordinates. Within the ipsilateral lateral area, we excluded saturated pixels and their adjacent regions up to 187.5 µm, similar to the local rolony exclusion method described earlier. Thresholding and morphological reconstruction were applied to remove autofluorescence backgrounds, including those from blood vessels. Additionally, we excluded the outer cortical edges of the flattened data (up to 212.5 μm on the cortical plate) due to distortions that occurred during the flattening process. To maintain consistency for comparative analysis, we also excluded regions extending beyond our dataset along the AP-axis. The criteria for defining medial and lateral areas are as shown in Fig. 4A. Furthermore, for consistency, we employed the same normalization factor (layer thickness) for normalized density as was used in the rolony analysis mentioned above.

To assess the differences in laminar projections within the samples from the Allen dataset, as well as between the Allen dataset and our current axonal BARseq dataset, we performed the two-sample Kolmogorov–Smirnov test (SupFig. 8C). In brief, the frequency distributions of projections along the cortical depth (SupFig. 8B) were transformed into cumulative distributions. We then calculated the Kolmogorov–Smirnov distances (KS-distances), using the maximum absolute difference between the two cumulative distributions. For comparisons within the Allen dataset (Allen vs. Allen), all-to-all comparisons were conducted for each cortical area, and the average KS-distance for each experiment was used to represent the differences within the dataset. For the comparisons between the Allen dataset and axonal BARseq data (Allen vs. axonal BARseq), we compared the bulk projections of rolonies (Fig. 4B) against individual experiments from the Allen dataset.

### Lat-projecting IT cell grouping

We selected Lat-projecting IT cells with a minimum of 5 rolonies in LatI/C and divided them into three groups based on soma layer (Fig. 5A). However, not all the barcoded cells have a known soma location (as described above), so we only included Lat-projecting cells with a known soma location in the analysis. As a control, we grouped barcoded cells without soma information into two groups based on projection depth, as shown in SupFig. 9G-I, M-N.

### Focal projection distance

For each barcoded cell, we calculated the all-to-all distances of the LatI/C rolonies on the ML-AP plate. For each rolony, we selected the shortest 33% of distances and used the mean of these distances to represent the ‘focal projection distance’ of the rolony. We then used the mean of the rolony focal projection distances to represent the distance for each cell.

We selected the shortest 33% distance as a measurement based on the following considerations: We wanted to capture the small range and tightness of the projection, so we calculated the focal projection distance using a subset of the nearest rolonies. We observed that cells can have more than one focal projection per target (e.g. the LatC projection from the orange cell in Fig. 5D), so we set a threshold (i.e. 33%) to exclude rolonies from other clusters. Based on our observations, most cells did not have many focal projections, and it was difficult to distinguish a cell with many focal projections from one with a sparse projection. Therefore, we assumed that cells have a maximum of three focal projections and calculated the distance from the closest 33% of rolonies.

To estimate the effect of rolony number on focal projection distance per cell, we randomly subsampled different numbers of rolonies from each barcoded cell and calculated the focal projection distance for each subsample. The ground truth distance was calculated using all rolonies from the same cell. We defined the sampling error as:

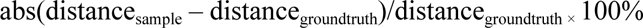

We performed 100 random samplings per cell, and the median error represents the error per cell. The results are shown in SupFig. 9J. Thus, we only used cells with ≥ 55 Lat rolonies for the following analysis (Fig. 5I, SupFig. 9K-N).

### Correlation between CT/CF fiber intensity and lower layer projection

Projections originating from deep-layer IT cells have been observed in the Allen Mouse Brain Connectivity Atlas (Oh et al., 2014), as indicated in SupFig 10A. Given that injections in WT mice can infect various cell types across depths, it’s important to estimate the proportion of infected deep-layer IT cells in individual experiments. Hence, we used CT/ET fiber intensity for estimation, based on the following considerations: when axons innervate a target, they often branch and establish multiple contacts, resulting in an increase in GFP intensity. However, this increase in GFP intensity might not be uniformly proportional across different cell types, injection sites, and animals. To minimize the influence of these variabilities, we chose to utilize the intensity of fibers before their innervation of subcortical targets. Furthermore, in this analysis, we assumed that the distribution of infected cells follows a unimodal pattern in the majority of experiments. This distribution comprises (1) layer 5 and above, (2) layers 5-6 and above, (3) layer 6.

We manually examined the fiber locations across 160 WT mice from the Allen Mouse Brain Connectivity Atlas. During this process, certain cortical regions were excluded primarily due to the following reasons: (1) CF routes entering the thalamus from the ventral side and dividing into smaller bundles to pass through the internal capsule (i.e., SSp-m, VISC). (2) The CT and ET routes following trajectories at angles that make their separation challenging when observed on a coronal plane (i.e., ACA, ORB). (3) A combination of these two conditions. Considering these factors, the validation process for the separation of CT/ET fiber bundles was conducted in the RSP, SSp-bfd/tr, AUD, and VIS areas.

In these regions, the dorsal and ventral routes of projections consistently aligned with the CT and ET fibers, as indicated in SupFig. 10B. To compare the intensity of the CT and ET fibers, we utilized the representation of the dorsal and ventral routes. The intensities of these two routes were quantified by manually drawing lines across the route on the stacks with the highest intensity. For each fiber, three lines were drawn, and the resulting mean intensity was calculated to represent the fiber’s intensity. To minimize the impact of saturation on measurements, we excluded mice with an injection volume exceeding 0.2 mm^3^. Additionally, to minimize potential bias, the evaluator was blinded to the cortical projection patterns. Mice with fiber bundles that were difficult to differentiate were excluded.

To maintain consistency with Fig. 5 and to minimize the effects of traveling fibers and intensity saturation near the injection site, we focused on measuring homotopic callosal projections. This approach involved manually defining a rectangular region of interest on the projection cluster situated closest to the mirror injection site on the flattened data of each experiment. However, this approach does introduce a potential concern that certain callosal projections may not terminate symmetrically in the contralateral area. For instance, callosal projections originating from the primary VIS cortex might not terminate in the contralateral primary VIS; they could be more likely to terminate at the boundaries of the primary VIS (Wang et al., 2011). To address this issue, we employed correlation analysis to filter out experiments lacking strong symmetric projection within the region of interest (Spearman’s rho < 0.5, along with manual confirmation). Additionally, samples with weak symmetric projections or injection sites positioned close to the flatmap’s edge were excluded from the analysis. The flattened cortical data were obtained as described above in the auditory GFP bulk tracing analysis. Because the regions of interest were selected manually, we did not include AP-axis data exclusion and autofluorescent background exclusion in this analysis. The normalized density was calculated using the layer thickness within the region of interest.

### Software and statistical analysis

We used MATLAB, ImageJ/FIJI, and MIJ (Sage et al., 2012) for data processing and visualization. We used MATLAB and GraphPad Prism 9/10 for statistical analyses, as indicated in the text. **, *p*-value < 0.01, ***, *p*-value < 0.001, ****, *p*-value < 0.0001, unless stated otherwise.

To estimate confidence intervals (2.5 and 97.5%), bootstrapping was performed for each group (2,000 iterations). * indicates that the confidence intervals of two groups have no overlap.

The data distributions presented in Figs. 4-5 and SupFigs. 8-9 are detailed in SupTable 5.

## Supporting information

SupFig. 1

SupFig. 2

SupFig. 3

SupFig. 4

SupFig. 5

SupFig. 6

SupFig. 7

SupFig. 8

SupFig. 9

SupFig. 10

SupFig. 11

SupTable 1

SupTable 2

SupTable 3

SupTable 4

SupTable 5

SupVideo

## Data availability

Raw and processed data will be available in the Brain Image Library: BIL uuid, dfabd518f37ff2e5 (Benninger et al., 2020). Code for image processing and analysis is available on GitHub (https://github.com/ZadorLaboratory/axonalBARseq).

## Acknowledgements

The authors would like to acknowledge Alex Vaughan and Justus M. Kebschull for VAMP2nλ library, Kathleen Lucere, Eugene Fong and John M. Bolger for virus injection and animal processing, Yu-Chi Sun, Wiktor Wadolowski and Barry Burbach for technical support, Longwen Huang, Daniel Fürth and Anand Suresh for useful discussion, chatGPT for manuscript editing, Allen Institute for Brain Science for the open access databases. This work was supported by the National Institutes of Health (RF1MH123403; U19NS123716) to A.M.Z; IARPA MICrONS (D16PC0008 to A.M.Z.), and Robert Lourie award (to A.M.Z.). In conducting research using animals, the investigator adheres to the laws of the United States and regulations of the Department of Agriculture.

## Author Contributions

L.Y., X.C., and A.M.Z. conceived the study. L.Y. performed experiments and data analysis. L.Y. and

A.M.Z. wrote the paper.

## Declaration of Interests

A.M.Z. is a founder and equity owner of Cajal Neuroscience and a member of its scientific advisory board.

## Supplementary tables

SupTable 1, List of brain sections and imaged areas SupTable 2, List of reagents and optics

SupTable 3, Experimental protocol modifications from BARseq

SupTable 4, Experiment numbers from the Allen Mouse Brain Connectivity Atlas SupTable 5, Data distributions of Fig. 4-5 and SupFig. 8-9.

## Supplementary video

SupVideo, Single-cell reconstruction from 100 barcoded neurons as shown in Fig. 2A.

---

**SupFig. 1.**
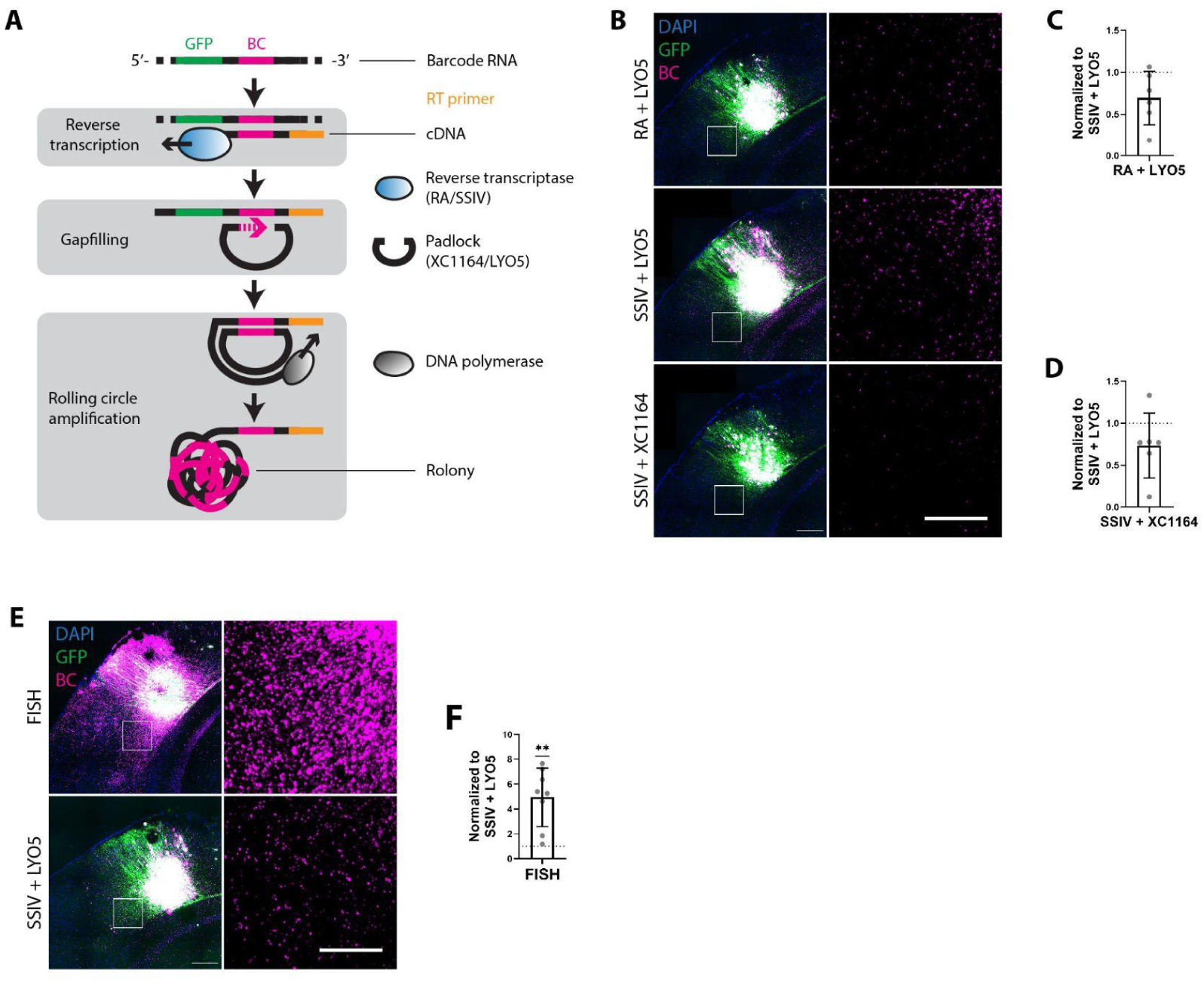
Optimization and efficiency estimation of axonal rolony preparation. (**A**) A brief summary of rolony preparation. Barcoded RNAs in axons and somata are first reverse transcribed (RT) into cDNA using reverse transcriptase. A padlock probe is then used to bind to the cDNA through adjacent regions of the barcode. The gap in the padlock is filled in using DNA polymerase, forming a circular single-stranded DNA molecule (ssDNA) through ligation by ligase. The circular ssDNA serves as a template for rolling circle amplification, resulting in the production of multiple copies of the barcode in a cluster of ssDNA called rolony. (**B-D**) Improvement of axonal rolony density using modified rolony preparation protocol. RA (RevertAid H Minus reverse transcriptase) and padlock probe XC1164 were used for reverse transcription and gap-filling for BaristaSeq (Chen et al., 2018). In the modified protocol, SSIV (SuperScript IV) and padlock probe LYO5 were used for reverse transcription and gap-filling. (B) Representative images of axonal rolonies using different rolony preparation conditions. Images were taken around the injection site in primary auditory cortex (AudI). Barcoded cells expressed both barcode RNA and GFP protein. (**C-D**) Both modifications yielded higher axonal rolonies density (one sample *t*-test). Each data point is the median of normalized rolony density from one brain section, two sections per mouse, total 3 mice. (**E-F**) The density of axonal barcodes was compared using two methods: FISH (RNAscope) and rolony preparation. Each barcoded RNA molecule has one GFP and one barcode. FISH detects the GFP component in the barcode RNA, while rolony preparation detects the barcode component. (E) Representative images of axonal barcode RNA around the injection site in AudI. (F) Comparison of the RNA density using FISH and the modified rolony preparation protocol (one sample *t*-test). Each data point is the median of normalized rolony density from one brain section, two sections per mouse, total 4 mice. Scale bar: top, 250 µm; bottom, 100 µm.

**SupFig. 2.**
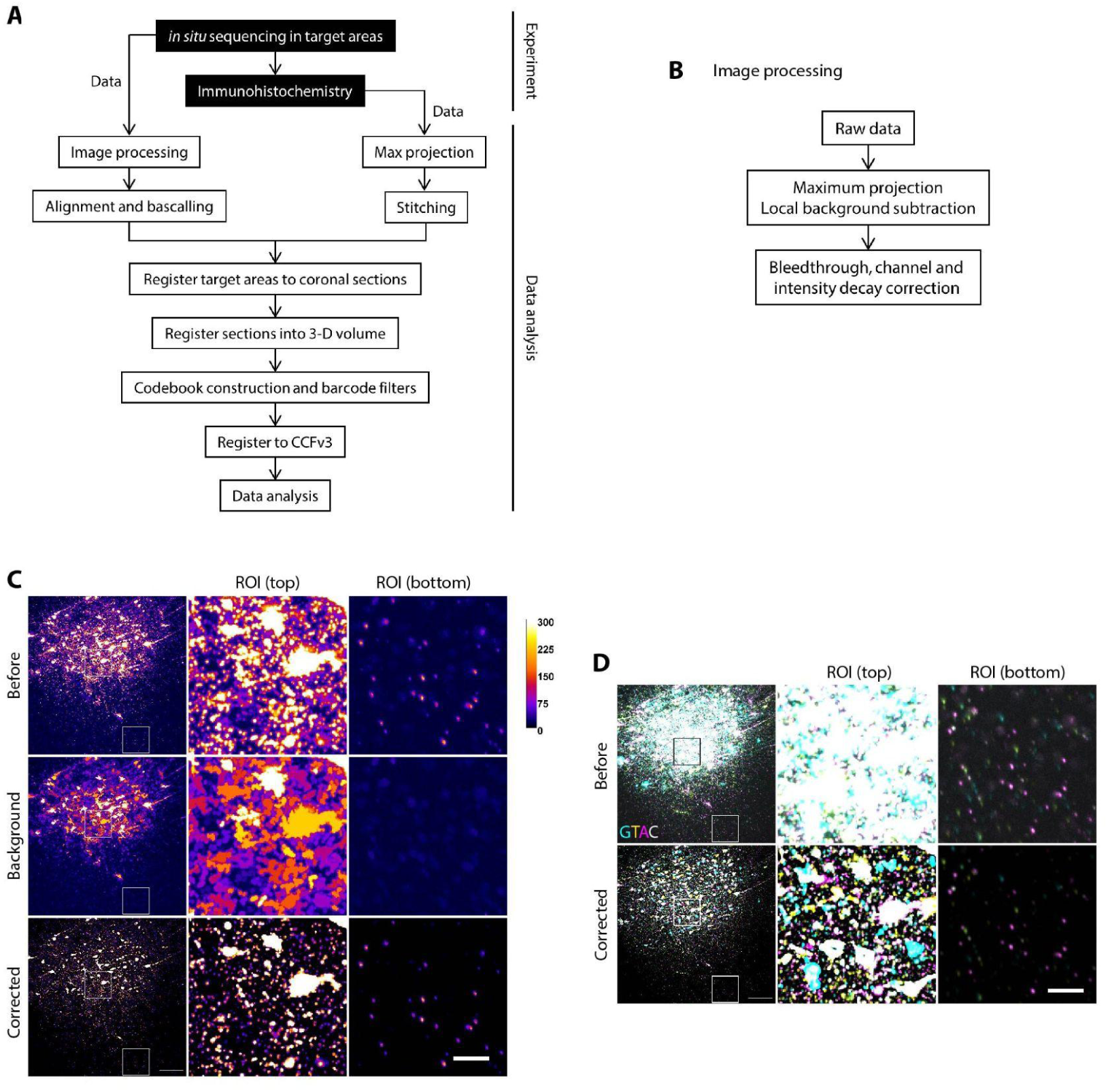
Data processing workflow and image processing. (**A**) The workflow of data acquisition and analysis for axonal BARseq. Additional information on image processing, alignment, base-calling, codebook construction, and barcode filtering can be found in SupFig 2-4. (**B**) The workflow of image processing. Raw data were acquired as z-stacks and then converted into maximum projections with background subtraction. Rolony intensity was then detected and used for further correction including bleedthrough, unequal channel intensity, and intensity decay during *in situ* sequencing. (**C-D**) Background subtraction effectively reduced the impact of uneven illumination, tissue and nucleus background on the images. The pixel intensity in the corrected images represented the signal beyond the background and was used for downstream analysis. Representative single tile from AudI, comparison of (C) single channel and (D) all four channels before and after background subtraction. Scale bar, 25 µm; color bar, single channel intensity.

**SupFig. 3.**
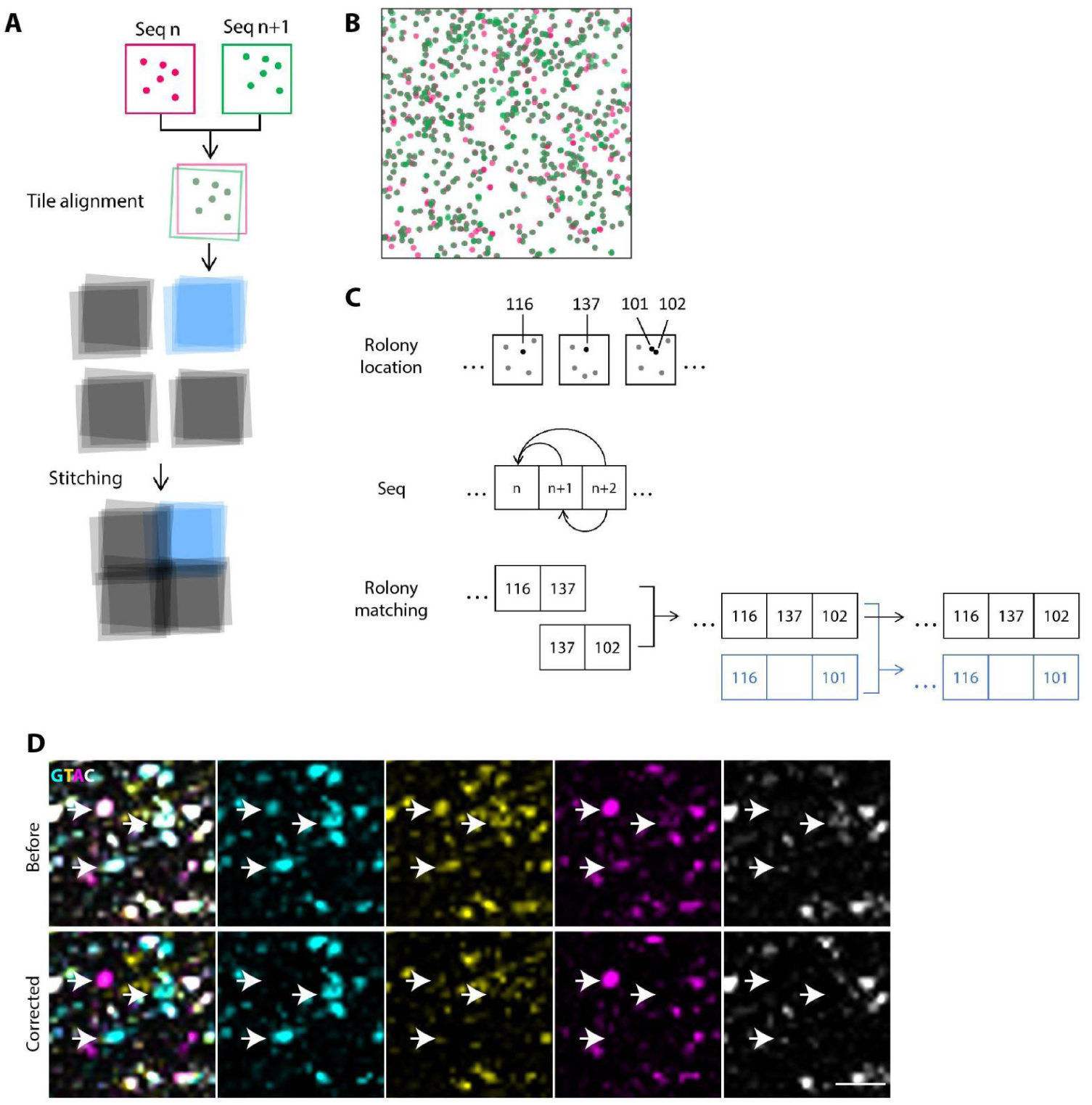
The processing steps for alignment, stitching and base-calling. (**A**) The workflow of alignment and stitching for axonal BARseq. Rolonies were identified as local maxima in individual tiles for each sequencing cycle. Tiles were first aligned for each cycle using the locations of the rolonies, and then the aligned tile stacks were stitched together using the locations of the rolonies in the overlap regions. (**B**) Example of point cloud registration between neighboring sequencing cycles (magenta and green). 400 × 400-pixel area from AudI. (**C**) The workflow of rolony base-calling. During each sequencing cycle, unique IDs were assigned to individual rolonies (top row). Rolonies were then matched to rolonies in multiple previous sequencing cycles (middle row). For each match, a pair of IDs was identified, and pairs from different matches were merged or branched into an additional sequence (blue). Barcode sequences were assembled using the nucleotide associated with each ID. If no ID was found in a particular cycle, no nucleotide was assigned and the grid remains empty. (**D**) Representative images for nonspecific signal correction in somata. Images were selected from Seq11 in AudI. Most of the somata had T in the previous sequencing cycle due to fixed barcode digits, and the signal was carried over to this cycle (top row). The correction effectively reduces the nonspecific signals (bottom row). Scale bar, 50 µm.

**SupFig. 4.**
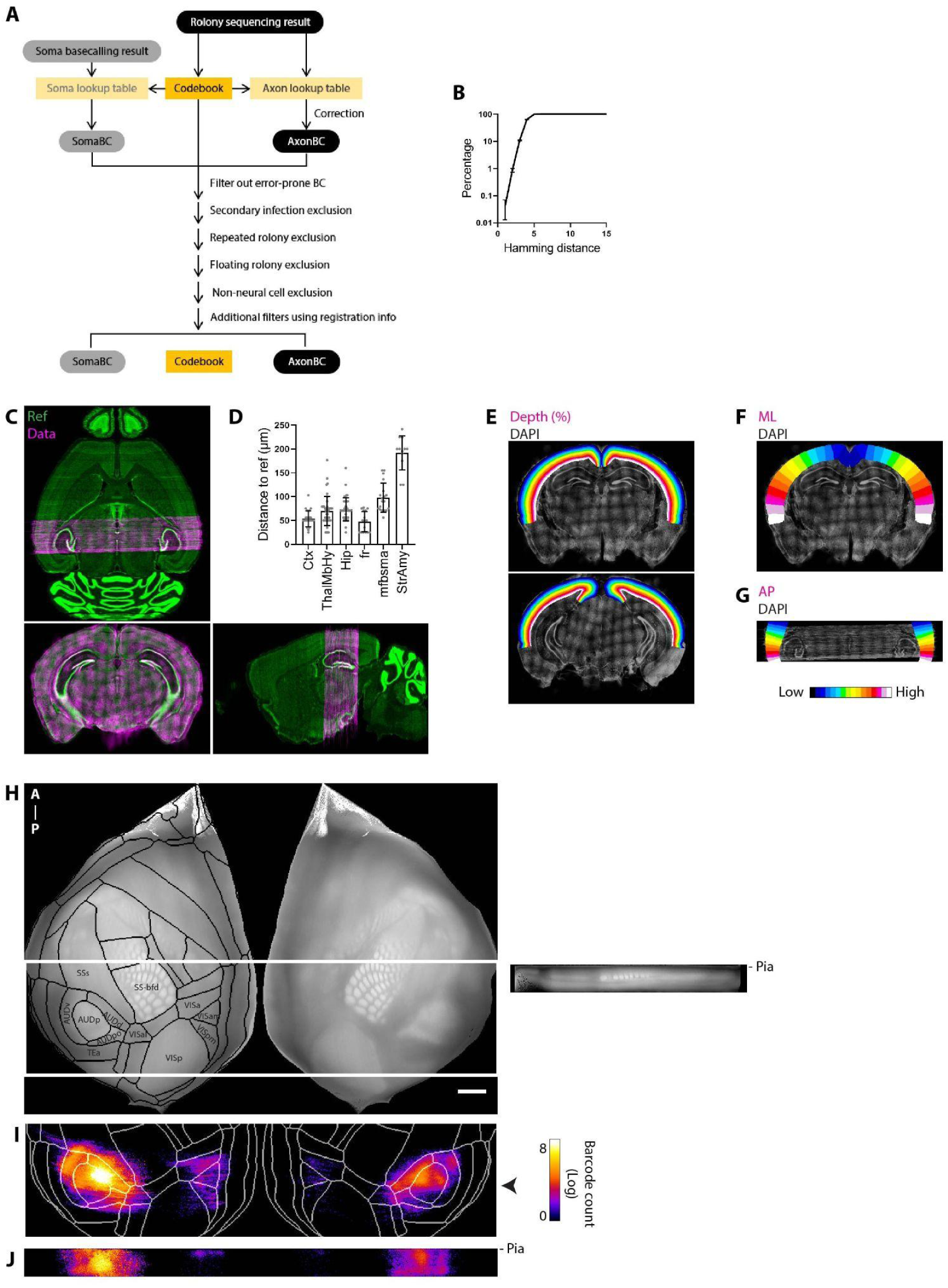
Barcode analysis and data registration. (**A**) The workflow of constructing a codebook and lookup tables for identifying somatic and axonal barcodes. (**B**) The possibility of barcodes having counterparts at various Hamming distances. 1figs4 15-nt random generated barcode, mean ± SD from 100 iterations. (**C**) Representative plates showing the registration of the data to the Allen Brain Atlas common coordinate framework (CCFv3). Magenta, DAPI staining from registered data; green, Nissl staining of the reference atlas. (**D**) Evaluation of CCFv3 registration. Distances between manual selections and CCFv3 boundaries (Mean ± SD). see *Methods*. Counts: Ctx, 24; ThalMbHy (midbrain/thalamus/hypothalamus), 42; Hip (hippocampal formation), 41; fr (fasciculus retroflexus), 16; mfbsma (mammillary related), 19; StrAmy (striatum/amygdala), 12. (**E-G**) Representative images of cortical depth/ML/AP-value of the registered brain. Colormaps encode continuous gradients for depth/ML/AP-axis. E-F, coronal view; G, horizontal view. Gray, DAPI staining from the data. (**H**) Cortical flatmap generated from CCFv3 average template. Grayscale: maximum intensity of the average template on ML-AP plate. White box: region of the axonal BARseq dataset. Area boundaries and annotation are shown in the left hemisphere. Area boundaries were drawn according to the most frequent area per pixel across cortical depth. Scale bar: 1 mm. Right panel, representative ML-depth view 125 µm slide of the right hemisphere. Top, cortical surface; bottom, the inner surface of the cortex. Depth percentage was converted to µm by assuming 1,000 µm cortical thickness. (**I-J**) Distribution of barcodes on the flatmap. (I) ML-AP view. White, area boundaries; arrow, AP-location of J. (J) Representative ML-depth view of barcode locations in cortex. Image included signals from 250 µm volume along the AP-axis.

**SupFig. 5.**
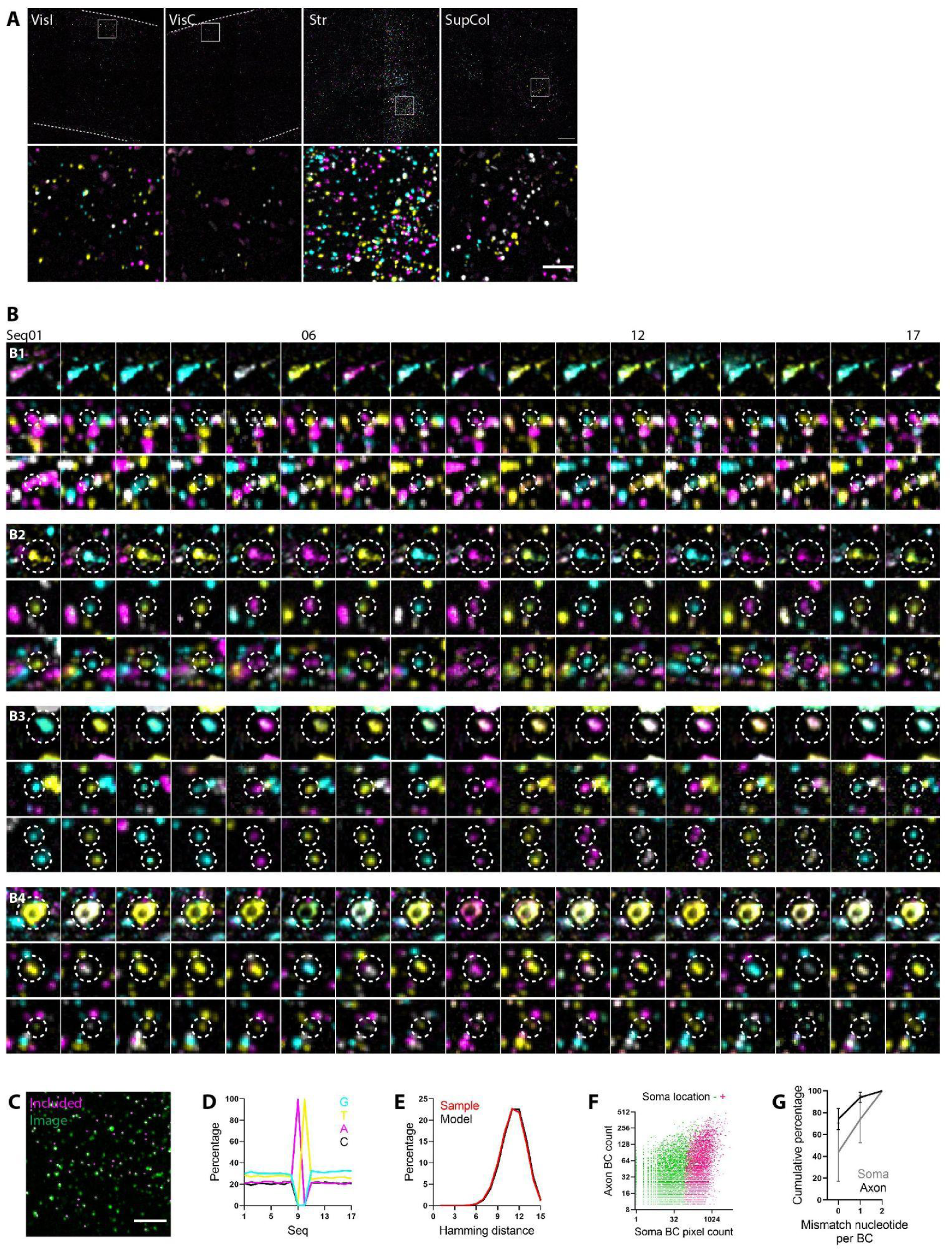
Example images and statistics of the dataset, related to Fig 1. (**A**) Representative images of axonal rolonies in VisI, VisC, Str and SupCol. Images are from Seq 01. Dotted line, cortical boundaries. Scale bar: top, 100 µm; bottom, 25 µm. Vis was identified according to CCFv3; it included posterior parietal cortex or parietal associative area (Lyamzin and Benucci, 2019). (**B**) Additional examples of somatic and axonal rolonies from the same barcode are shown to illustrate the variability of sequencing signals in this dataset. Four examples are presented, with the top row showing images from somata and the middle and bottom rows showing images from axonal rolonies. The examples display differences in the size, shape, and intensity of axonal and somatic rolonies. Some somata exhibit phasing (B3-4; phasing correction is described in SupFig. 3D), while others have clean signals (B2). In B2 and B3, duplications of the same rolonies were found in some sequencing cycles due to imperfect stitching, which was corrected in a later processing step (see *Methods*). (**C**) Example of the proportion of barcoded rolonies included in this dataset. Rolonies included in this dataset are indicated in magenta. Scale bar, 25 µm. (**D**) Nucleotide proportion of barcodes across sequencing cycles. The 9-10th nucleotides were fixed for this barcode library, 99.1% of the barcodes contained both fixed nucleotides. (**E**) The distribution of the all-to-all Hamming distance of the sample was similar to that of the model. Sample, 15-nt barcode without the two fixed nucleotides. Model, 10,000 random generated 15-nt barcodes, mean ± SD of 100 iterations. (**F**) The relationship between the counts of somatic and axonal barcodes in individual barcoded cells. Cells were divided into two groups based on whether or not their soma location was identified. Cells that were somatic location-negative may still have barcodes detected in AudI, but their somatic locations were excluded from the analysis due to potential errors (see *Methods*). (**G**) Cumulative percentage of mismatch nucleotides per barcode in axon and soma. Barcodes with soma location were used to compute the soma barcode percentage; mean ± SD. Because the current codebook was constructed using axonal barcodes, it resulted in higher accuracy in identifying barcodes in axon compared to soma. VisC/I, contra/ipsilateral visual cortex; Str, striatum.

**SupFig. 6.**
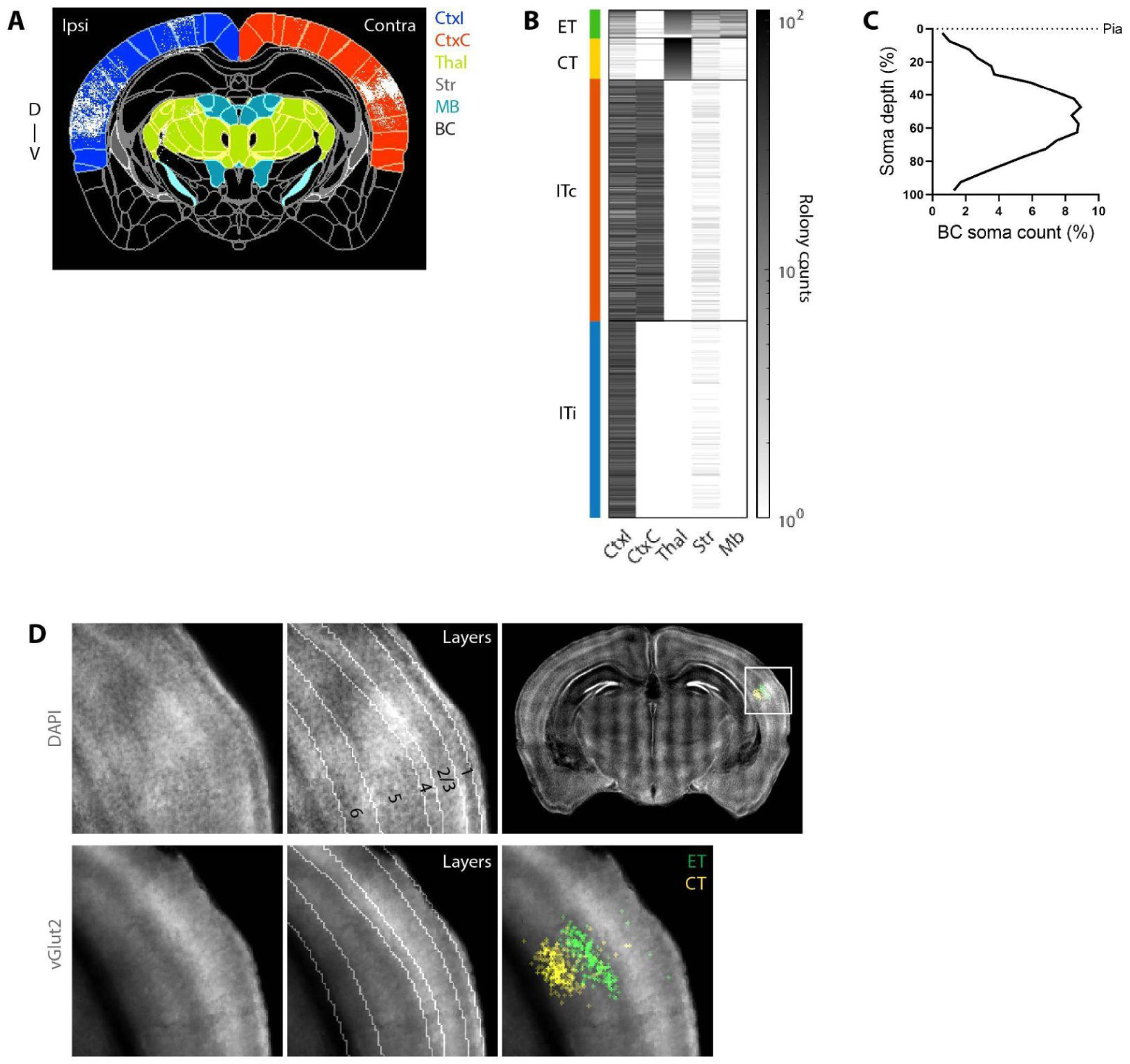
Projection targets of major cell types, related to Fig 3. (**A**) Representative coronal view of rolonies and five targeted brain regions. Rolonies located within the fiber bundles were not included in the five regions, 468026 of 492950 rolonies were included. Target regions are color-coded; gray line: anatomical boundaries. (**B**) Heatmap of rolony counts in different targets from four cell types, ET, CT, ITc and ITi. (**C**) Distribution of barcoded somata along the cortical depth. Total 3,698 barcoded somata. Bin size: 5% cortical depth. (**D**) Coronal view of the layer boundaries and internal markers. Zoom-in view: injection site. Top row: DAPI; bottom row: vGlut2. Consistent with previous studies, layer 4 has relatively strong vGlut2 and DAPI signals (Nahmani and Erisir, 2005; Coleman et al., 2010; Sato et al., 2022). Conversely, the vGlut2 signal is relatively weak in upper layer 5 (Hirai et al., 2012; Oswald et al., 2013). The Images were generated from a 312.5 µm volume of the registered data. Soma positions are sum projections and markers are median projections of the volume. MB, midbrain.

**SupFig. 7.**
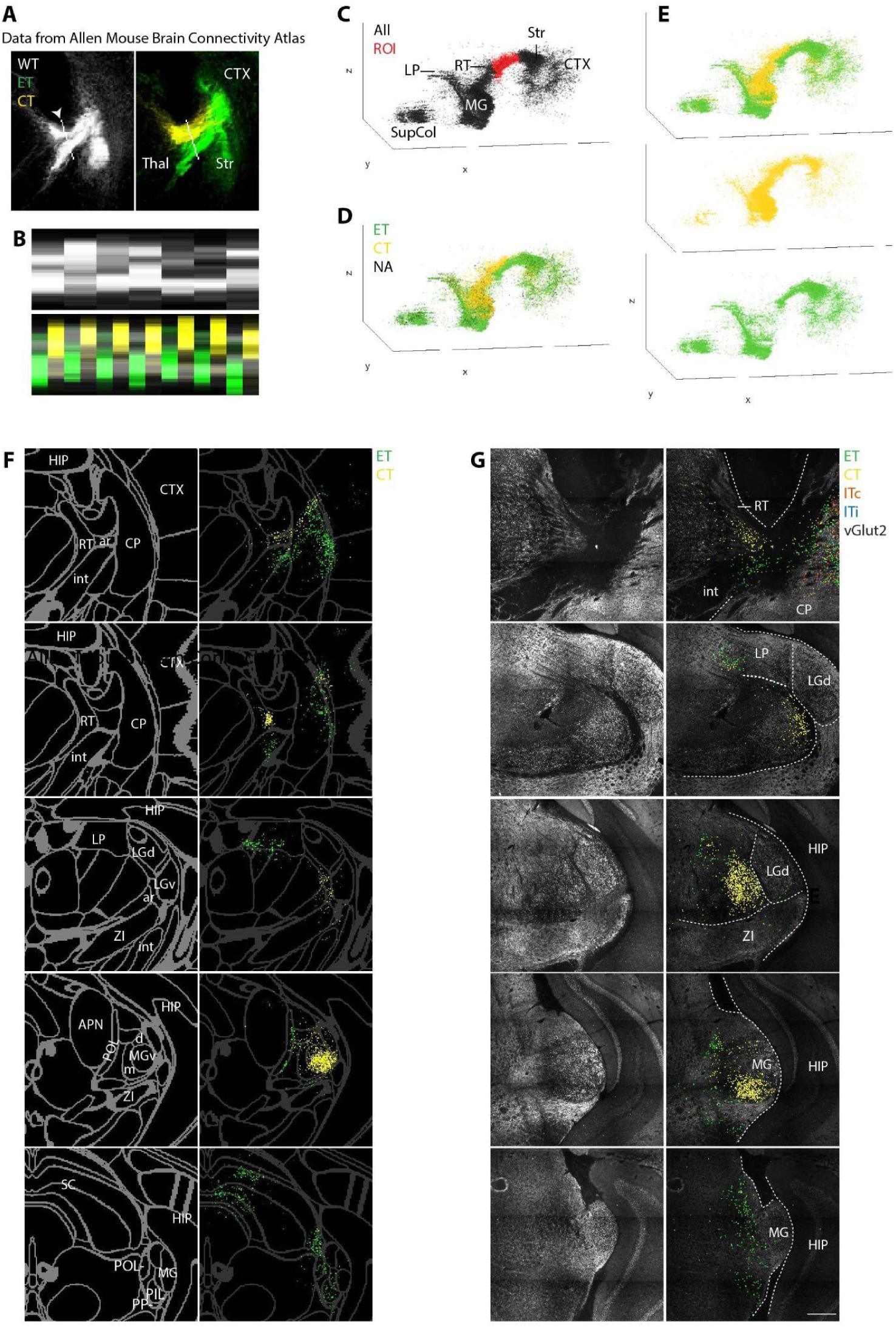
Division of CT and ET cells using projection targets and axonal trajectory, related to Fig 3. (**A-B**) The dorsal and ventral routes to the thalamus of CF cells in the Allen Mouse Brain Connectivity Atlas. In bulk tracing experiments conducted on AUD, two distinct trajectories were observed in wild-type mice (WT, C57BL/6J and Emx-Cre). Conversely, the CT and ET Cre lines (CT: Ntsr1 and Syt6; ET: Rbp4 and Sim) predominantly displayed the dorsal and ventral routes, respectively. (A) The coronal view illustrates the locations of the trajectory routes between the striatum and thalamus in WT, CT, and ET mice. The white line (*arrow*) indicates the line used for intensity profiling in (B). Panel (B) provides intensity profiling of the two routes on a single plate from WT, CT, and ET mice. Each white column represents a WT mouse, while the corresponding locations of the CT and ET routes are shown below for each mouse. The length of the white line and intensity of projections were normalized for visualization purposes. Reference-aligned images were utilized with a voxel size of 25 µm. CT and ET intensity values were combined from multiple Cre mice to represent the projections from AUD. Experiment number: SupTable 4. (**C-E**) The region of interest (ROI) and results from CT/ET grouping of Thal+ cells. (C) Side view of rolonies from Thal+ cells (black). Red, rolonies within the ROI (around RT and striatal-thalamic fiber tract), used for grouping CT and ET for Thal+ cells. (D) Initial grouping result using rolony within the ROI. Group identities were assigned to barcoded neurons according to their rolony locations within the ROI; individual rolonies were color-coded based on their cell identity. Black dots, rolonies from cells without rolony in ROI. (E) Final grouping result. Cells without rolony in the ROI were assigned to the group as their closest neighbors (see *Methods*). All Thal+ cells were assigned to either ET or CT. (**F**) Localization of registered CT and ET rolonies in striatum, thalamus and superior colliculus. Left column, region boundaries with labels; right column, rolonies in the same region with tune-down boundaries for visualization. Each row is one coronal section, arranged from anterior (top) to posterior (bottom), from 6,925, 7,050, 7,500, 8,150 and 8,775 µm in CCFv3, 25 µm/section. (**G**) Localization of CT and ET rolony with vGlut2 as an internal marker for thalamic structures. Left column, vGlut2 staining; right column, rolonies in the same region with tune-down vGlut2 signal for visualization. MG contains large but lower density of vGlut2 puncta (Hackett et al., 2016). To preserve the original resolution of vGlut2 staining, images and rolony locations were not registered. Each row is one coronal section, arranged from anterior (top) to posterior (bottom), from 220, 720, 1,120, 1,560 and 2,020 µm in this dataset, 20 µm/section. Scale bar: 250 µm. CTX, cortex; CP, caudoputamen; ar, auditory radiation; RT, reticular nucleus; int, internal capsule/cerebral peduncle; HIP, hippocampus; LP, lateral posterior nucleus; LG(d/v), lateral geniculate complex (dorsal/ventral part); APN, anterior pretectal nucleus; POL, posterior limiting nucleus; MG(v/d/m), medial geniculate complex (ventral/dorsal/medial part); PIL, posterior intralaminar thalamic nucleus; PP, peripeduncular nucleus; ZI, zona incerta; SC, superior colliculus.

**SupFig. 8.**
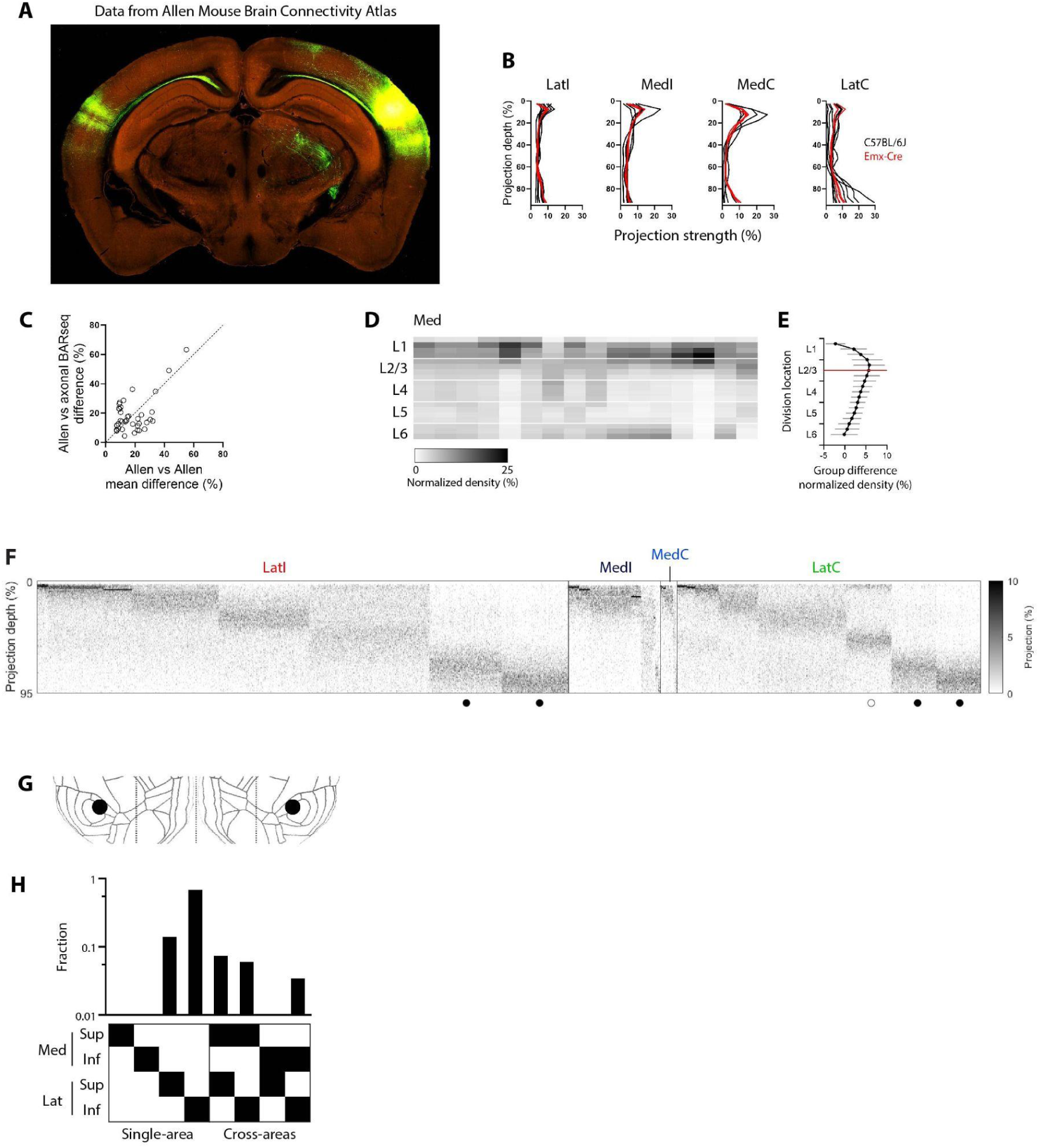
Description and comparison of laminar patterns, related to Fig 4. (**A-E**) AUD bulk tracing results and related analyses in wild-type mice (C57BL/6J and Emx1-IRES-Cre) from Allen Mouse Brain Connectivity Atlas. (A) Representative image. Experiment number, 158314278. (B) The GFP intensity profiling in LatI/C and MedI/C along cortical depth. Bin size, 5% depth. (**C**) The bulk tracing results from axonal BARseq are similar to those of the Allen dataset. We computed the differences in projection distributions across the four cortical areas, both within the Allen dataset and between the Allen dataset and axonal BARseq data. The Kolmogorov–Smirnov distance (KS-distance) was used to represent the differences in projections (see *Methods*). The results across the four areas were combined. The differences are distributed symmetrically on both sides of the diagonal line. Additionally, all KS-distances between the Allen and axonal BARseq datasets fall below the one-tailed 95% confidence interval of the KS-distances within the Allen dataset. (D) The normalized density in MedI/C along cortical layers. Each layer is divided into four bins along depth to visualize changes within the layer. Each column represents one Med area from a mouse. The deepest bin was excluded due to insufficient data points. (E) We divided the Med-projection into two groups using the mid-L2/3 as the dividing point. To determine the optimal division for the Med-projection, we divided it into two groups at various positions along the layers (y-axis). For each division, we calculated the difference in mean projection density above and below the division (x-axis). The divisions spanning from the bottom 1/4 of L1 to the top 1/2 of L2/3 exhibited the maximum group difference. Thus, mid-L2/3 was selected as the division point for the Sup and Inf groups. Mean ± SD. B-E, all target areas were manually checked and areas with few projections were excluded (see SupTable 4). (**F**) Heatmap of single-cell projection strength along the depth of the corresponding cells in Fig. 4D. Bin size, 1% depth. (**G-H**) The fractions of superficial and inferior projections to different areas in LatC local-exclusion control (with rolonies in the mirror somatic region of LatC excluded). (G) The exclusion area in LatC is represented by an additional black disk. (H) Quantification as Fig. 4F.

**SupFig. 9.**
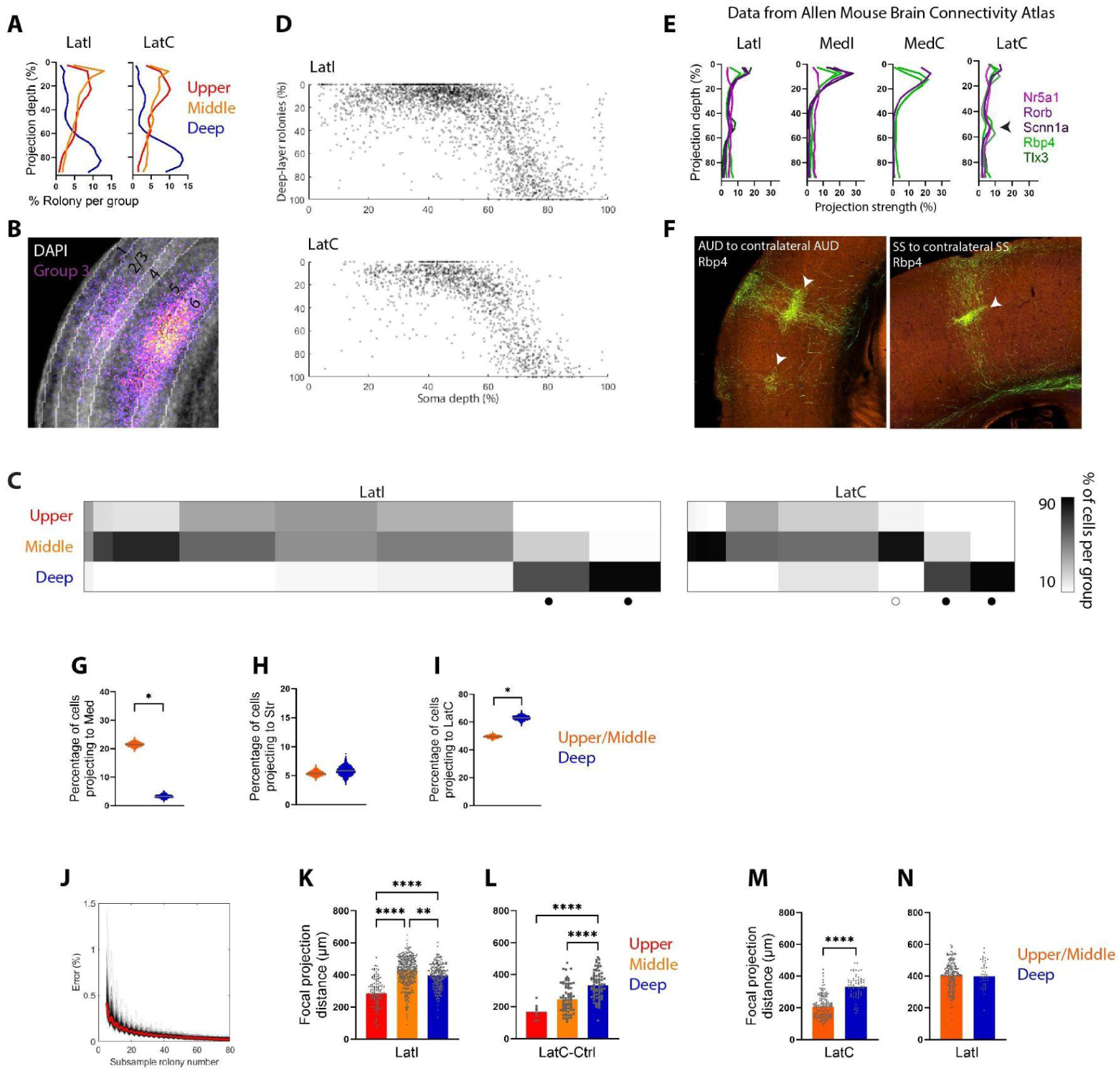
Projection differences across IT cell groups, related to Fig 5. (**A-B**) The bulk projection profiling of barcoded cells across cortical areas. (A) At the bulk level, deep Lat-projecting somata mainly projected to deep layers while upper/middle somata projected to all layers with preference to upper-middle layers. The deep-layer projections from deep somata extend from lower layer 5 to layer 6 in LatC (B). (**C**) The percentage of cells in each of the three groups corresponds to the clusters shown in Fig. 4D. The width of each cluster is as shown in Fig. 4D, and the percentage of cells was calculated for each respective cluster. Cells without soma location information were excluded. (**D**) At the single-cell level in the Lat areas, a gradual increase in deep-layer projection is observed along the depth of the soma. Each dot represents a single cell, with a minimum of 10 rolonies per cell. Cell counts: LatI, 2,723; LatC, 1,459. Among deep somata, there is a correlation between soma depth (%) and deep-layer rolonies (%) for both LatI and LatC; Spearman’s correlation, *p*-value (two-tailed) ****; cell counts: LatI, 733; LatC, 429. (**E-F**) The laminar projection profiling across cortical areas in layer 4 and 5 IT Cre lines with auditory injections. Notably, mid-layer projections were observed in contralateral Lat areas (indicated by arrow), frequently accompanied by layer 1 projections. (F) These images represent mid-layer terminations of homotopic callosal projections originating from Rbp4-Cre mice. It’s worth noting that this pattern is not limited to the auditory cortex and can be observed in other regions as well. Experiment number: AUD, 182090318; SS, 297652799. (A, E) Lines represent the frequency distribution of rolony depth/GFP intensity per group. Bin size, 5% depth. E, all target areas were manually checked and areas with few projections were excluded (see SupTable 4). (**G-I**) Projection target difference between upper/middle and deep layer-projecting cells without soma location information. Lat-projecting cells were categorized into upper/middle and deep layer-projecting cells based on their median rolony depth in LatI/C: upper/middle, layer 1 to upper layer 5; deep, lower layer 5 to layer 6. In 1,958 biLat cells, 198 (10.1%) were assigned to different groups in two Lat regions and they were excluded. Cell counts: upper/middle, 3,106; deep, 923. Consistently, upper/middle layer-projecting cells had a higher percentage of cells projected to Med (G) and a lower percentage of cells projected to LatC (I). The lack of difference observed in (H) could be attributed to the fact that cells projecting to the upper/middle layers consist of both upper and middle layer neurons. Gray line, median; G and I, *, significant difference with no overlap CI. (**J**) Effect of rolony number on computing focal projection distance. LatC-projecting cells with minimum 80 rolonies were used for testing, and 5-80 rolonies were randomly sampled per cell. The error caused by downsampling was calculated using the distance difference between downsampling and ground truth (see *Methods*). Ground truth distance was represented by distance computed with all rolonies. Black line, median error from each barcoded cell, total 314 cells; solid red line, median of black lines. The median error < 5% when sample rolony number ≥ 55, therefore, the minimum rolony number for this test was set to 55. (**K-L**) Projections from upper somata were more focal compared to deep somata in LatI (K) and in LatC local-exclusion control (L). Bar graph, median; dots, individual cells. Cell counts: LatI, upper, 122; middle, 326; deep, 211. LatC-Ctrl, upper, 8; middle, 72; deep, 77. Kruskal-Wallis test, Dunn’s test for multiple comparisons. (**M-N**) Projections from cells projecting to the upper/middle layers were more focal compared to those projecting to the deep layers in the LatC (M). The grouping of upper/middle and deep layer-projecting cells is as described in (G-I). The lack of difference observed in (N) could be attributed to the fact that cells projecting to the upper/middle layers consist of both upper and middle layer neurons. Mann-Whitney test; cell counts: LatI, upper/middle, 181; deep, 43. LatC, upper/middle, 145; deep, 73. SS, somatosensory areas.

**SupFig. 10.**
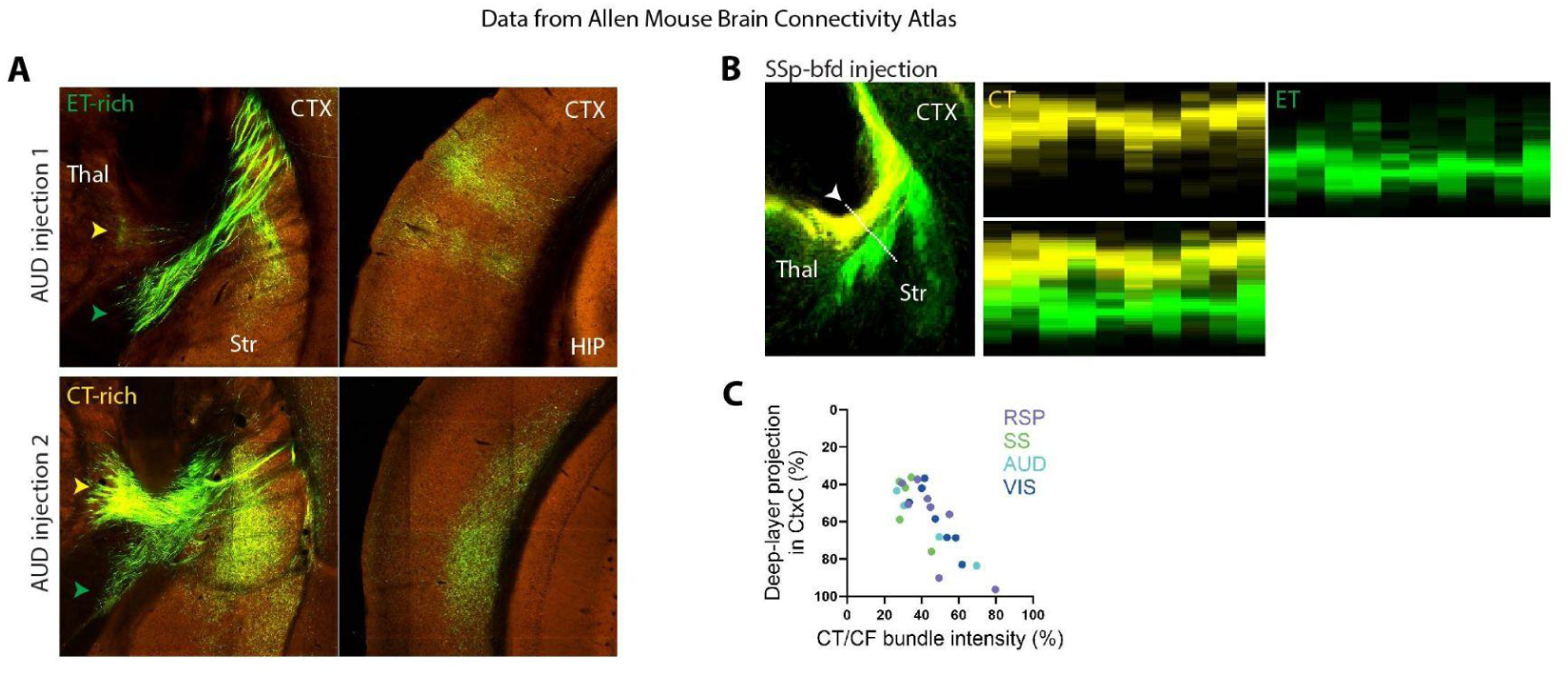
Indirect evidence obtained by comparing injection-variability across bulk tracing experiments of WT mice from Allen Mouse Brain Connectivity Atlas supports the claim that callosal homotypic projections of deep-layer neurons preferentially terminate in deep layers of the target. (**A**) We compared two AUD injections. The injection (*top left)* preferentially labeled more superficial (non-CT) neurons, as indicated by the very weak projection to reticular nucleus (*yellow arrow*) compared with a dense labeling of the internal capsule (*green arrow*), whereas the injection (*bottom left*) labeled a higher number of deep thalamic-projecting CT neurons. We use the differences in the labeling of these two pathways as a surrogate for the fraction of deep vs upper-middle labeled neurons. The corresponding contralateral projection targets showed that more superficial injections had more superficial contralateral targets (*top right*) compared with the targets of deeper injections (*bottom right*). Experiment number: top, 139519496; bottom, 112881858. (**B**) The distinction between the two CF projection routes is consistently observed across various cortical areas (see *Methods*). We compared the locations of CT and ET routes using CT and ET-Cre mice, as described in SupFig. 7A-B. One column represents injection in one cortical area. From left to right: AUDp, SSp-tr, SSp-bfd, RSPagl, RSPd, RSPv, VISal, VISam, VISl, VISp. CT and ET intensity were combined from one/multiple Cre mice to represent the projections from the cortical areas. Cre lines: ET, Rbp4 and Sim1; CT: Ntsr1 and Syt6. (**C**) Correlation between the proportions of deep-layer projection and CT/CF fiber bundle intensity. Spearman’s correlation, *p*-value (two-tailed) ****; experiment counts: RSP, 8; SS, 5; AUD, 4; VIS, 7. CF bundle intensity was represented as the sum of the intensities of CT and ET bundles. Deep-layer projection was defined as projections in lower layer 5 and layer 6 as Fig. 5. Experiment number: SupTable 4. All the data were reference-aligned images with a resolution of 25 µm/voxel. SSp-tr/bfd, primary somatosensory area, trunk/barrel field; AUDp, primary auditory area; RSPagl/d/v, retrosplenial area, lateral agranular/dorsal/ventral part; VISal/am/l/p, anterolateral/anteromedial/lateral/primary visual area.

**SupFig. 11.**
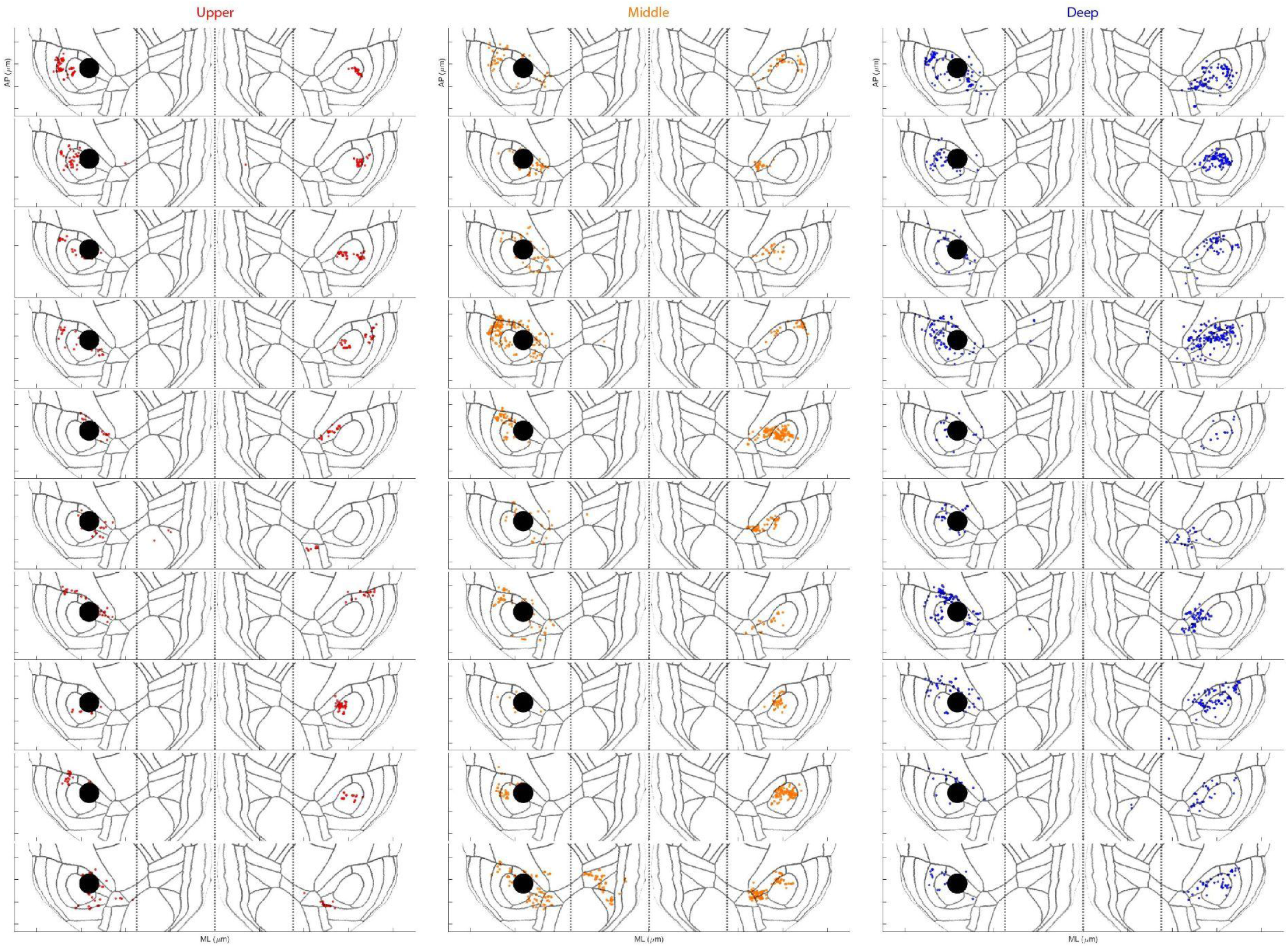
Additional examples of rolony location of individual neurons on cortical flatmap, as shown in Fig. 5D.

## Notes

### Summary of Updates

Revision of Analysis in Fig. 4: we have substantially revised the analysis to study the diversity of laminar terminations from intratelencephalic neurons across various cortical areas. Validation with Allen Mouse Brain Connectivity Atlas: we have added multiple panels to compare our results with the bulk tracing results from the Allen Mouse Brain Connectivity Atlas. Enhanced Description of Cortical Projection: To strengthen the analysis clarity, we have revised the manuscript to describe cortical projections using layer distribution along with cortical depth.

