## Supplementary figures and images for "Massive Multiplexing of Spatially Resolved Single Neuron Projections with Axonal BARseq"

### SupFig. 1

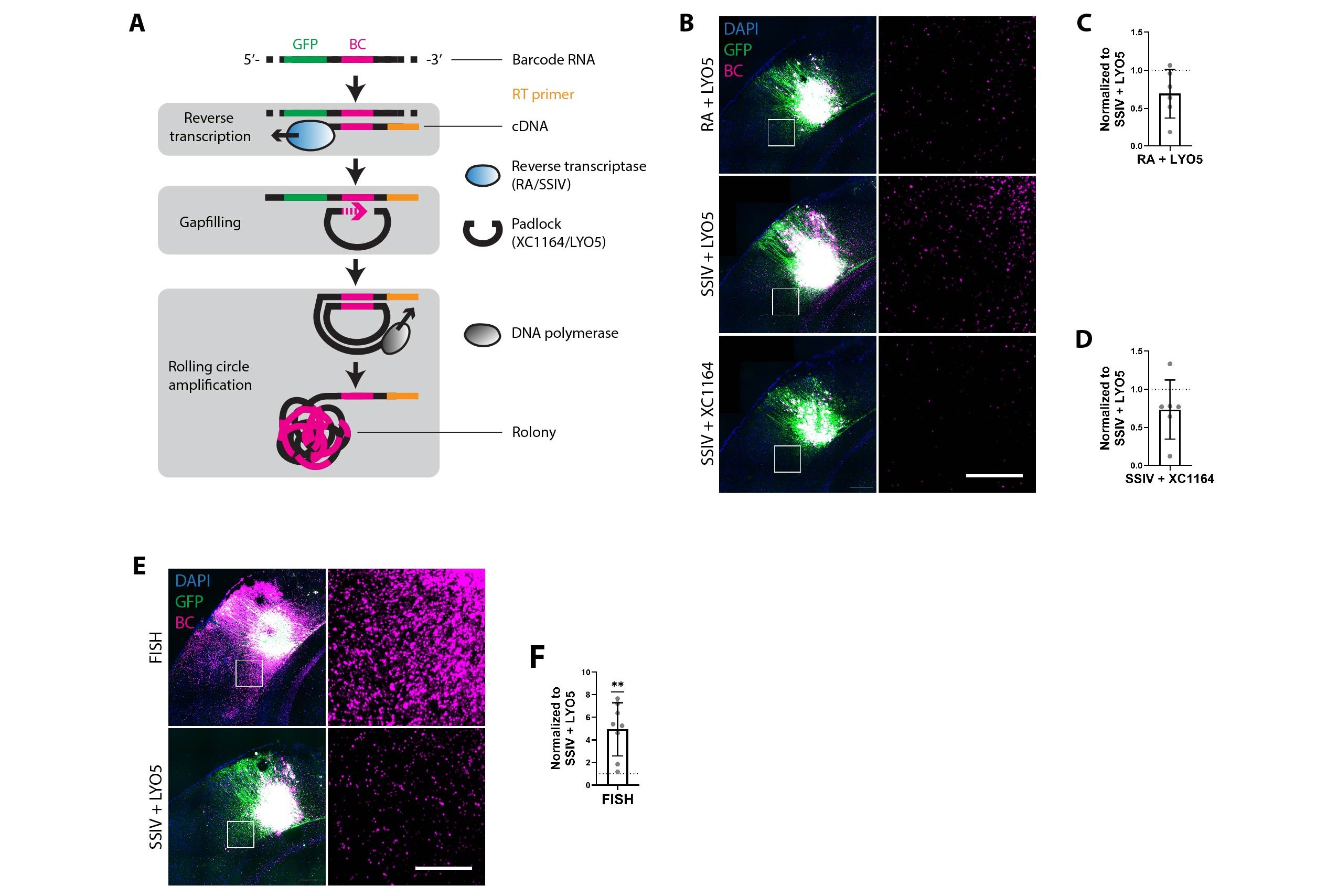

### SupFig. 2

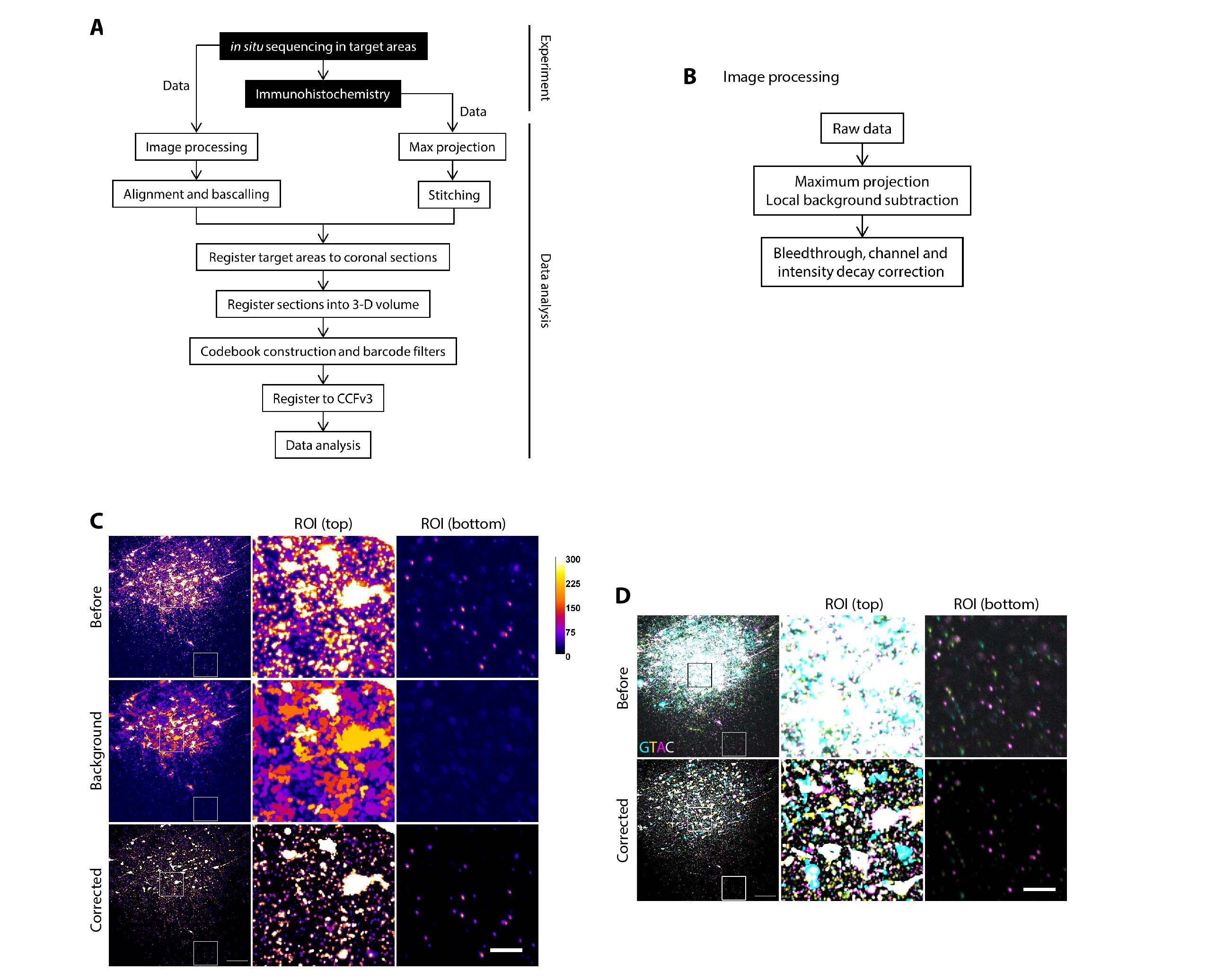

### SupFig. 3

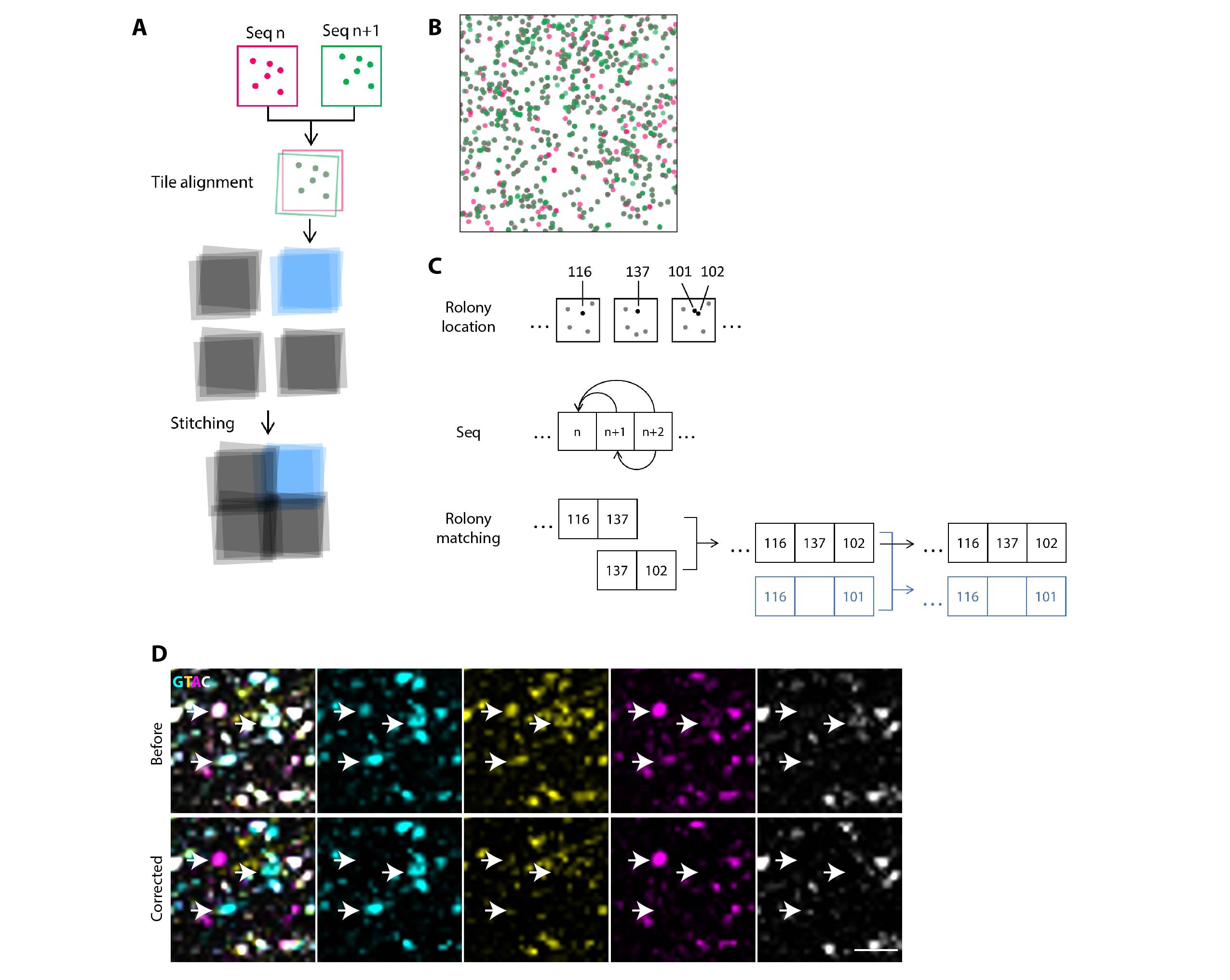

### SupFig. 4

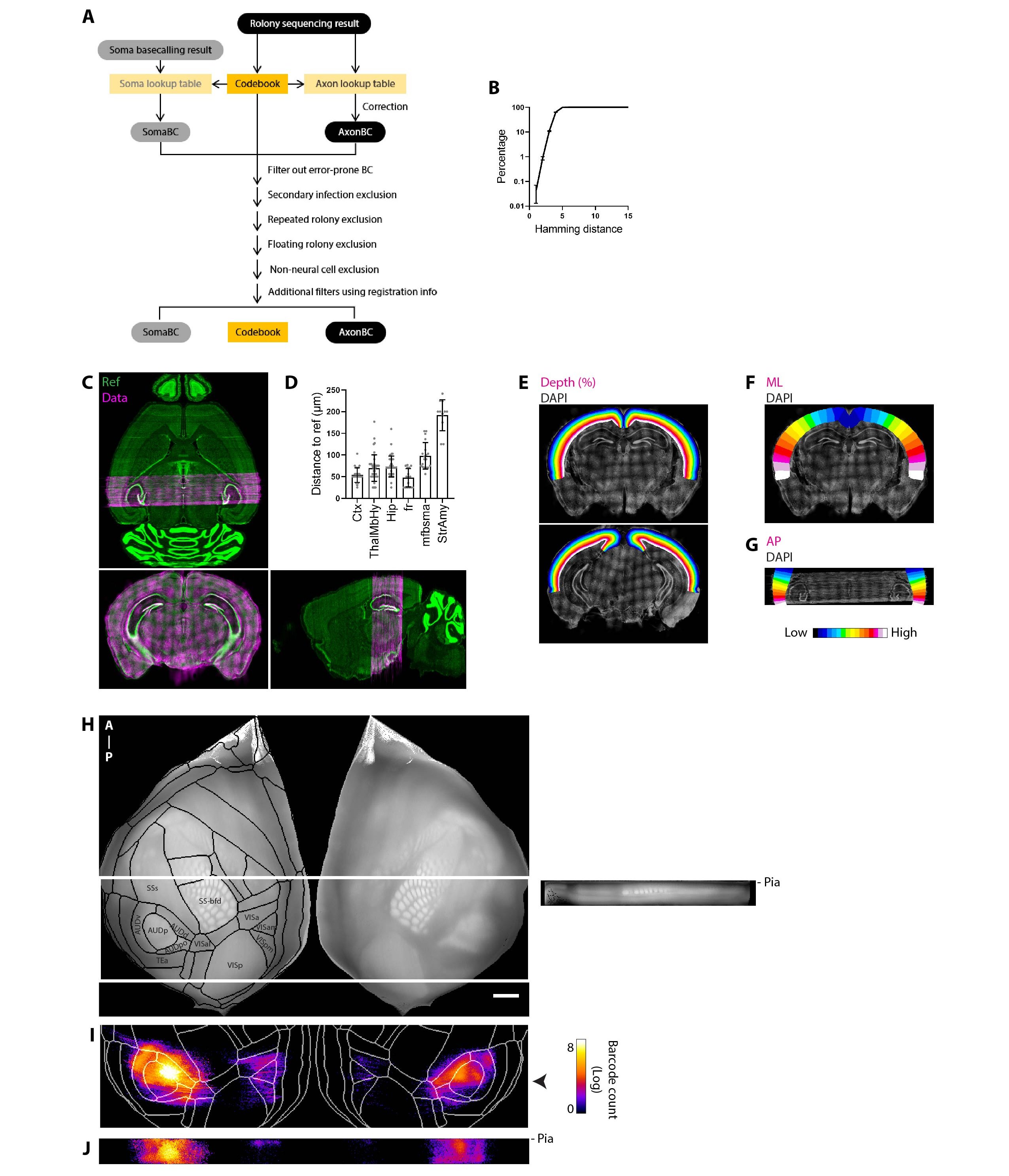

### SupFig. 5

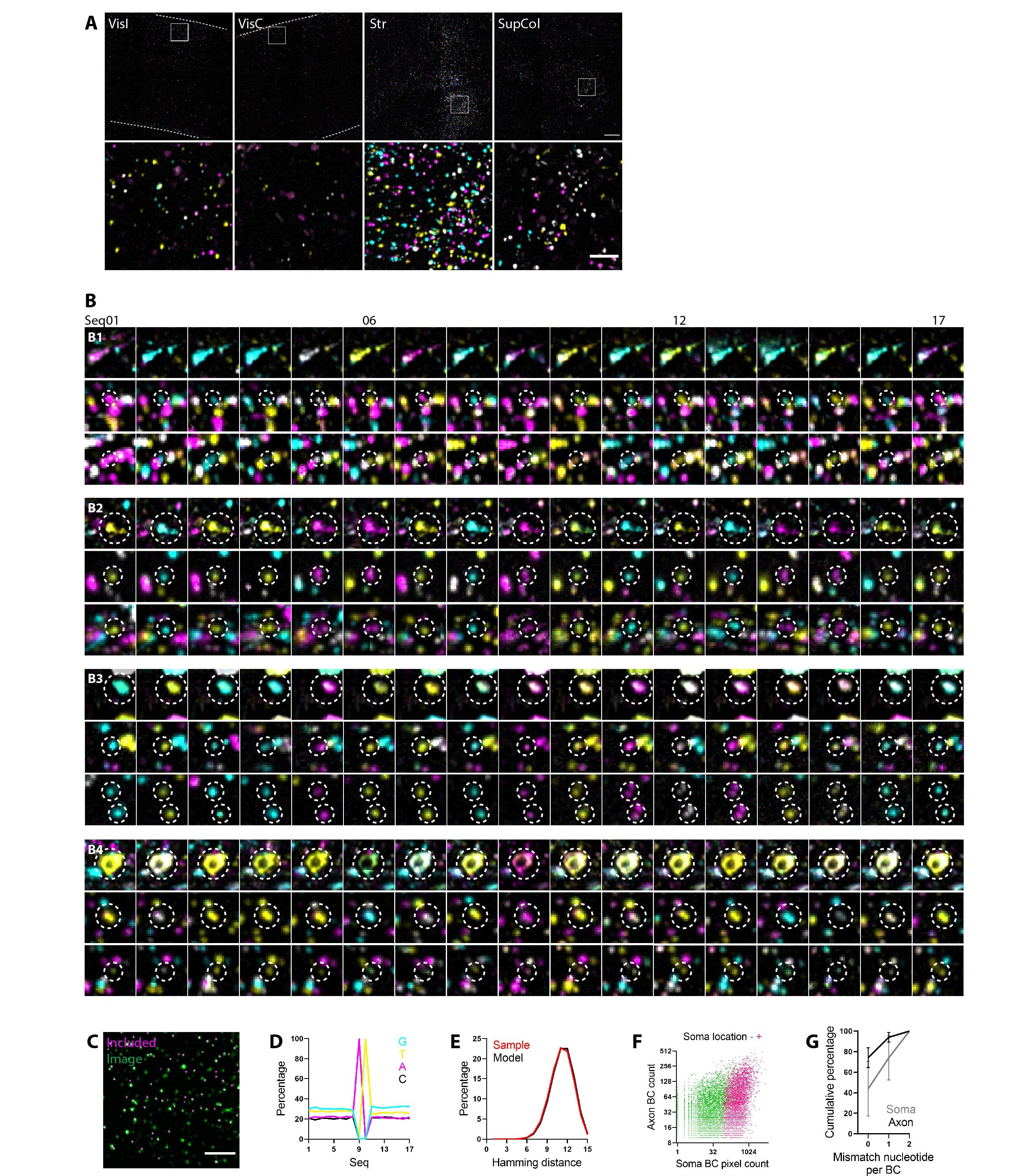

### SupFig. 6

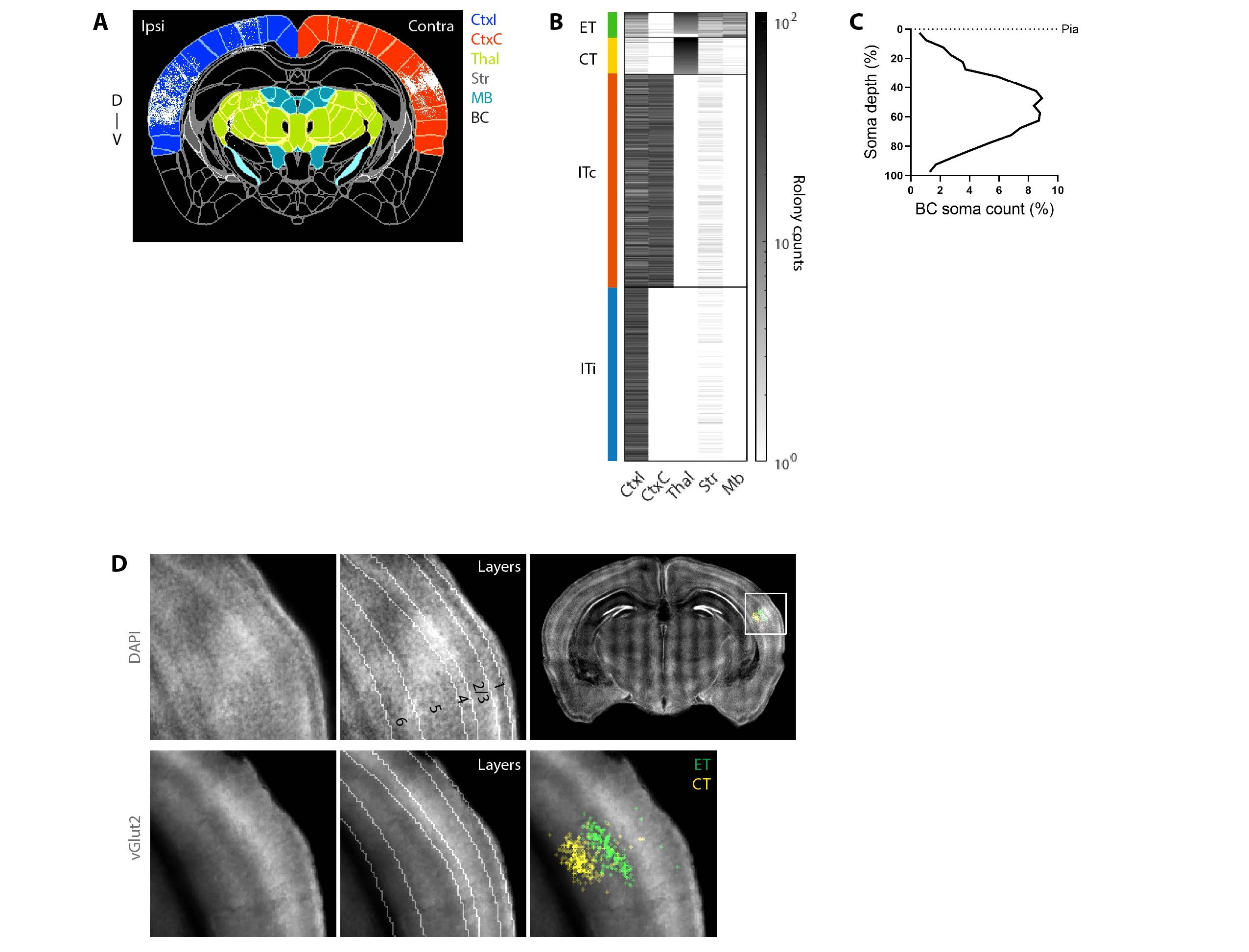

### SupFig. 7

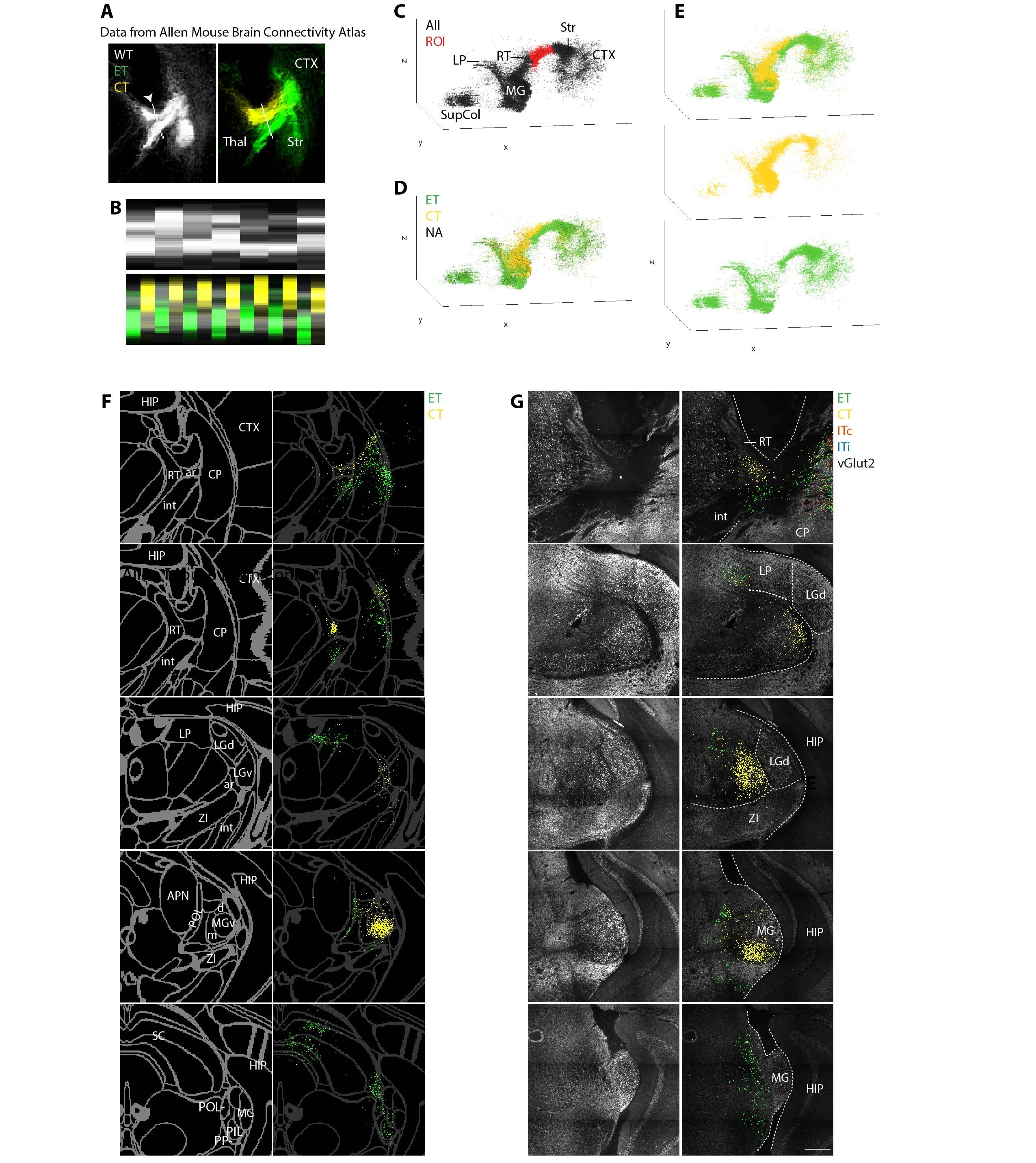

### SupFig. 8

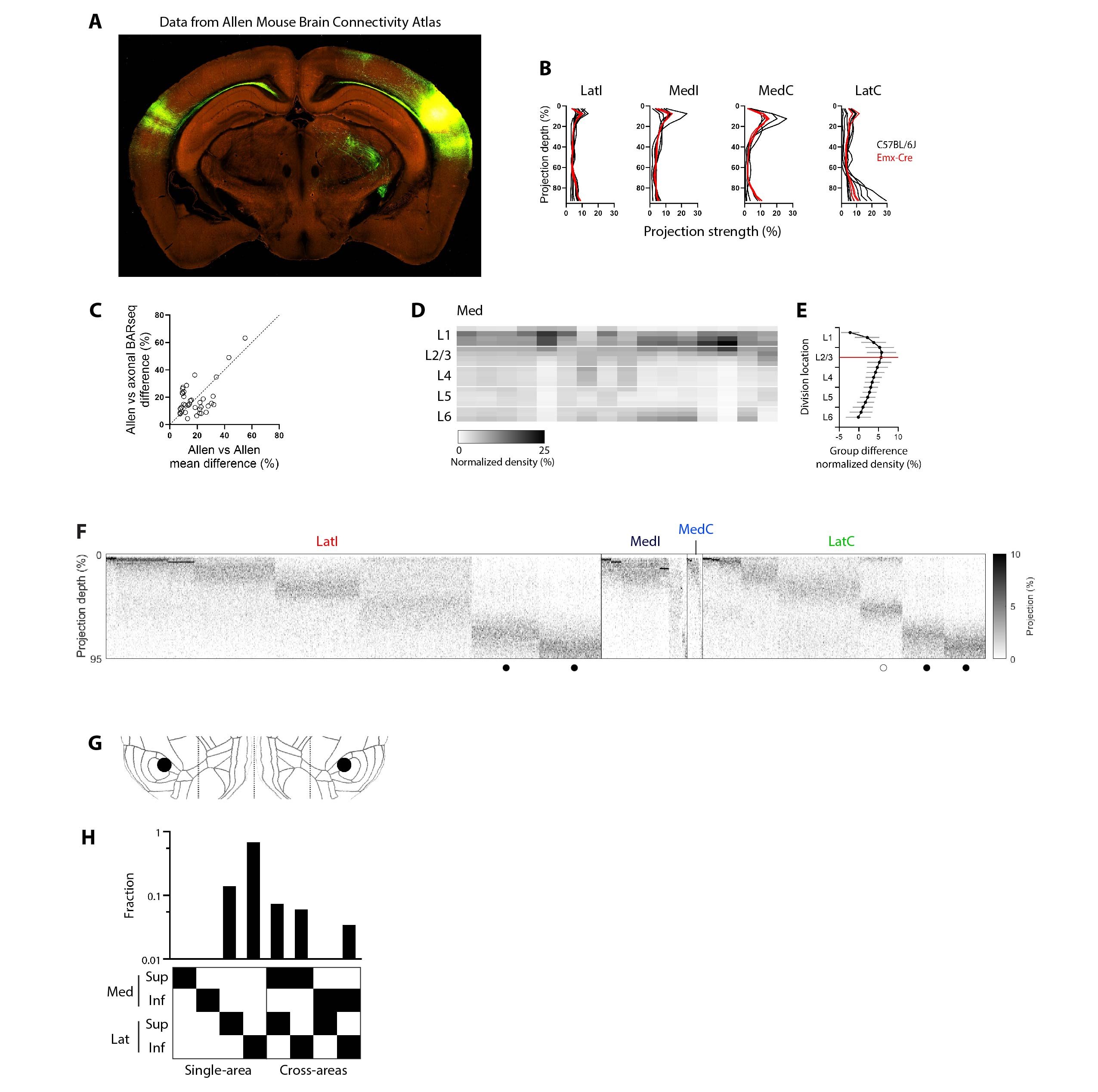

### SupFig. 9

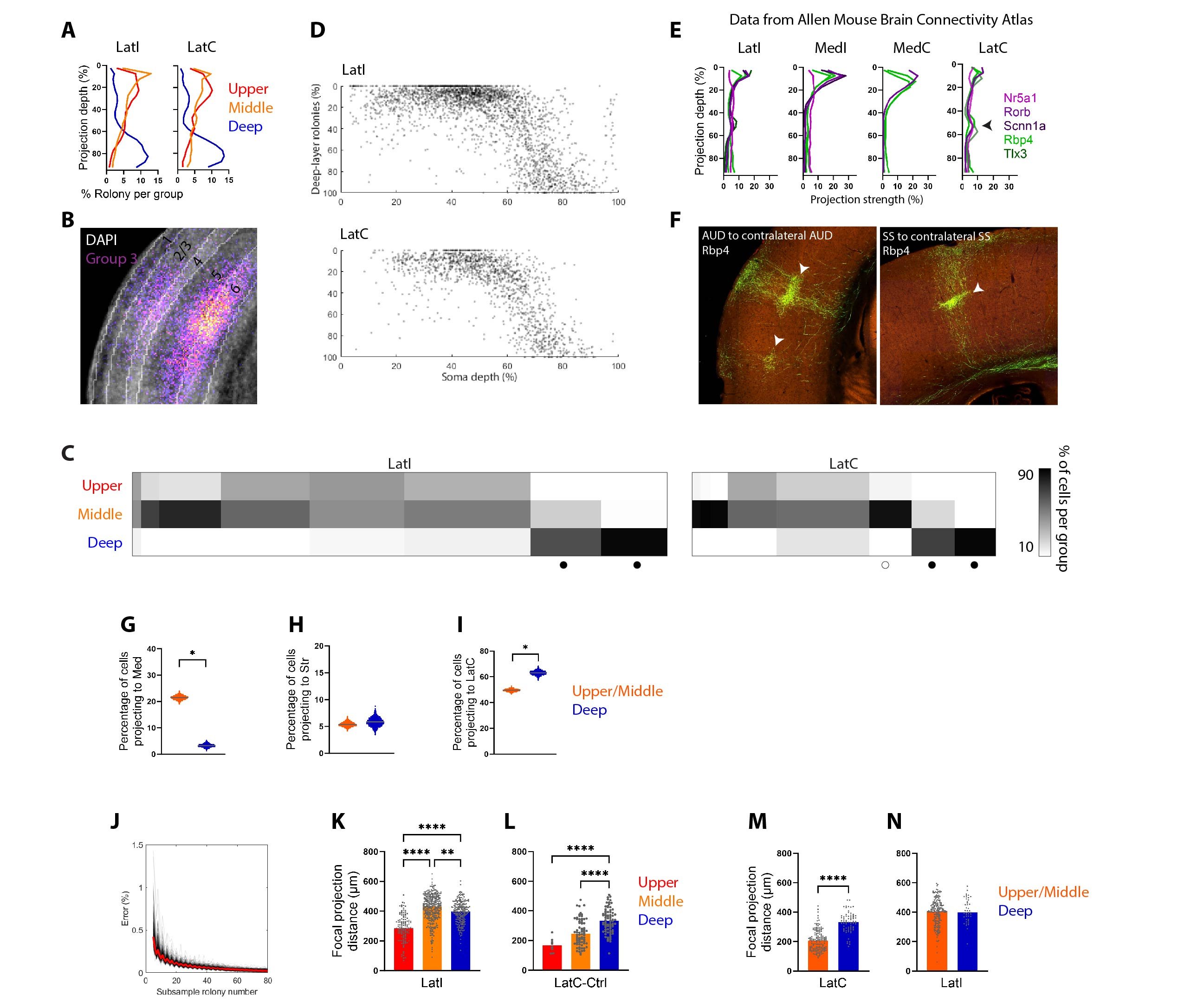

### SupFig. 10

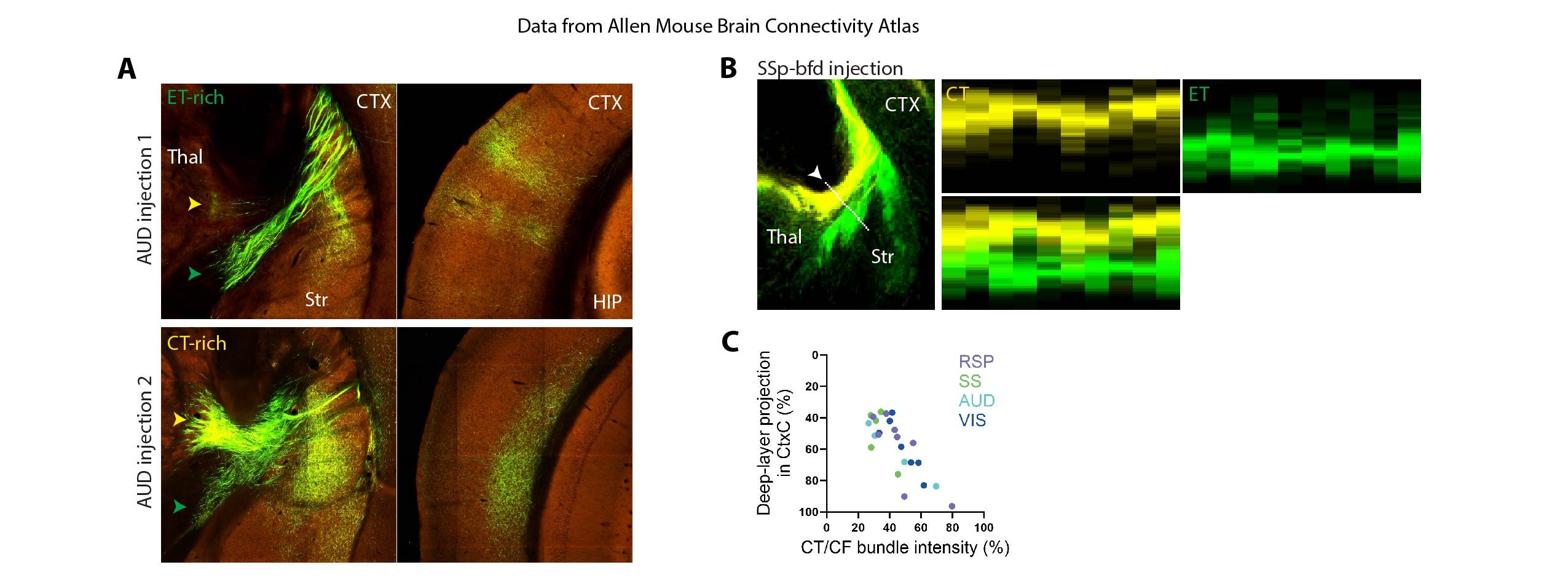

### SupFig. 11

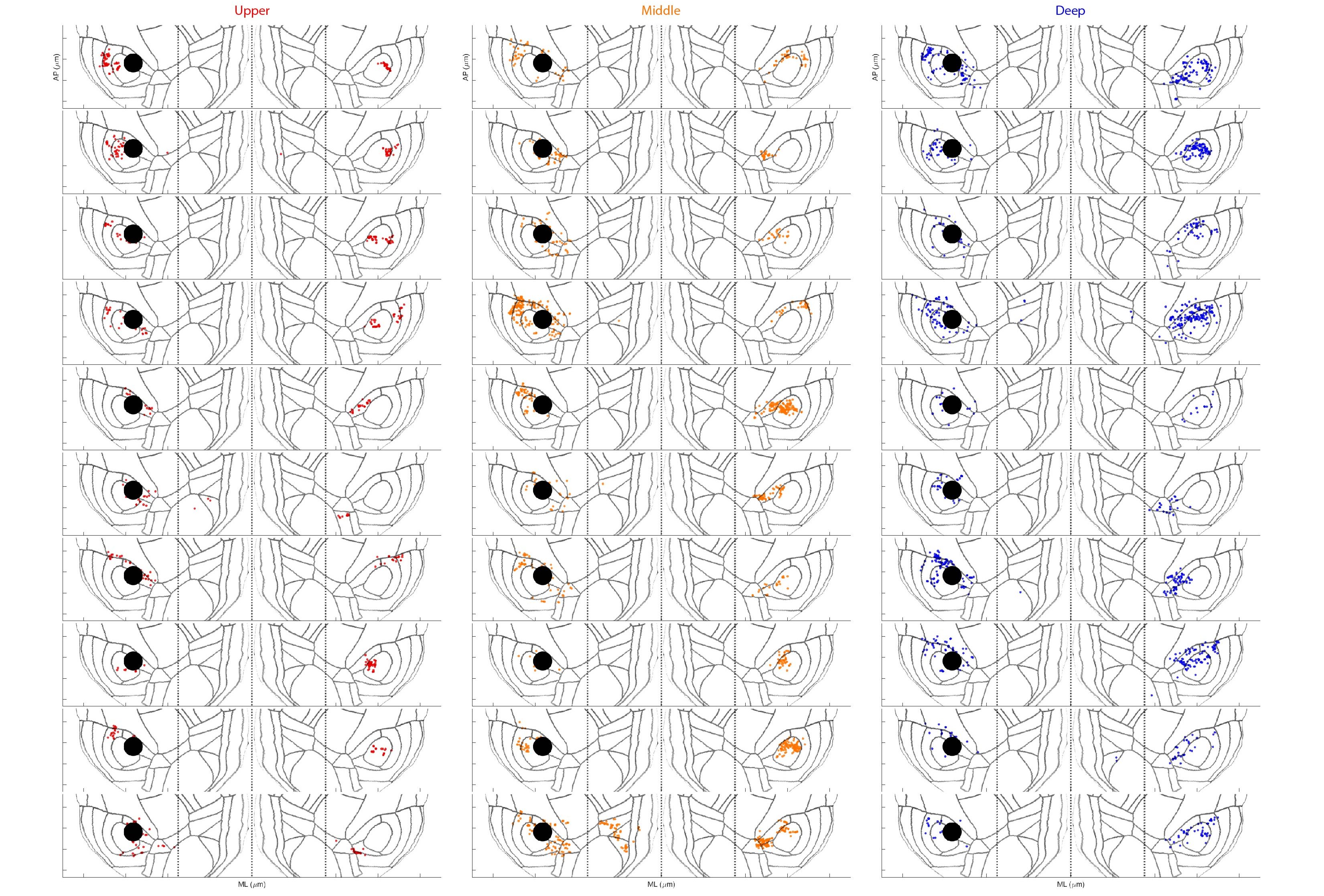
